# An adaptive teosinte *mexicana* introgression modulates phosphatidylcholine levels and is associated with maize flowering time

**DOI:** 10.1101/2021.01.25.426574

**Authors:** Allison C Barnes, Fausto Rodríguez-Zapata, Karla A Blöcher-Juárez, Daniel J Gates, Garrett M Janzen, Andi Kur, Li Wang, Sarah E Jensen, Juan M Estévez-Palmas, Taylor M Crow, Heli S Kavi, Hannah D Pil, Ruthie L Stokes, Kevan T Knizner, Maria R Aguilar-Rangel, Edgar Demesa-Arévalo, Tara Skopelitis, Sergio Pérez-Limón, Whitney L Stutts, Peter Thompson, Yu-Chun Chiu, David Jackson, David C Muddiman, Oliver Fiehn, Daniel Runcie, Edward S Buckler, Jeffrey Ross-Ibarra, Matthew B Hufford, Ruairidh JH Sawers, Rubén Rellán-Álvarez

## Abstract

Native Americans domesticated maize (*Zea mays* ssp. *mays*) from lowland teosinte *parviglumis* (*Zea mays* ssp.*parviglumis*) in the warm Mexican southwest and brought it to the highlands of México and South America where it was exposed to lower temperatures that imposed strong selection on flowering time. Phospholipids are important metabolites in plant responses to low-temperature and phosphorus availability, and have been suggested to influence flowering time. Here, we combined linkage mapping with genome scans to identify *High PhosphatidylCholine 1* (*HPC1*), a gene that encodes a phospholipase A1 enzyme, as a major driver of phospholipid variation in highland maize. Common garden experiments demonstrated strong genotype-by-environment interactions associated with variation at *HPC1*, with the highland *HPC1* allele leading to higher fitness in highlands, possibly by hastening flowering. The highland maize *HPC1* variant resulted in impaired function of the encoded protein due to a polymorphism in a highly conserved sequence. A meta-analysis across HPC1 orthologs indicated a strong association between the identity of the amino acid at this position and optimal growth in prokaryotes. Mutagenesis of *HPC1* via genome editing validated its role in regulating phospholipid metabolism. Finally, we showed that the highland *HPC1* allele entered cultivated maize by introgression from the wild highland teosinte *Zea mays* ssp. *mexicana* and has been maintained in maize breeding lines from the Northern US, Canada and Europe. Thus, *HPC1* introgressed from teosinte *mexicana* underlies a large metabolic QTL that modulates phosphatidylcholine levels and has an adaptive effect at least in part via induction of early flowering time.

## Introduction

Elevation gradients are associated with changes in environmental factors that impose substantial physiological constraints on an organism. Adaptation to highland environments is achieved via the selection of genetic variants that improve their ability to withstand lower oxygen availability (1; 2), increased ultraviolet (UV) radiation (3) and lower temperatures (4). In particular, cold temperatures reduce thermal time accumulation, measured in growing degree days (GDDs), (5) and select for accelerated development and maturity as a compensatory mechanism (6). Following domestication from teosinte *parviglumis* (*Zea mays* ssp. *parviglumis*) (7) in the lowland, subtropical environment of the Balsas River basin (Guerrero, México), cultivated maize (*Zea mays* ssp. *mays*) expanded throughout México and reached the highland valleys of central México around 6,500 years ago (8).

In México, highland adaptation of maize was aided by substantial adaptive introgression from a second teosinte subspecies, teosinte *mexicana* (*Zea mays* ssp. *mexicana*), that had already adapted to the highlands of México thousands of years after its divergence from teosinte *parviglumis* (9; 10). Adaptation to low temperature and soils with low phosphorus content in highland environments drove *mexicana* genetic divergence from the lowland *parviglumis* (11). Phenotypically, the most evident signs of *mexicana* introgression into maize are the high levels of stem pigmentation and pubescence (12) that are thought to protect against high UV radiation and low temperatures. The ability to withstand low temperatures and efficiently photosynthesize during the early stages of seedling development are key factors in maize highland adaptation (13). Indeed, recent transcriptome deep sequencing (RNA-seq) analysis showed that the inversion *Inv4m*, introgressed from *mexicana*, strongly affects the expression of genes involved in chloroplast physiology and photosynthesis (14). Given the slow accumulation of GDDs in typical highland environments, selection has favored shorter generation times in highland adapted maize (15). By the time maize reached the Mexican highlands, its range had already expanded far to the South, including the colonization of highland environments in the Andes (16; 17). Andean maize adaptation occurred largely independently of *mexicana* introgression (18; 19), and there is no known wild teosinte relative in South America. These multiple events of maize adaptation to highland environments make maize a good system to study the evolutionary and physiological mechanisms of convergent adaptation (18; 19).

In comparison to its southward expansion, the northward migration of maize into the modern-day United States, where summer daylengths are longer, occurred at a much slower pace (20; 21) due to delayed flowering of photoperiod-sensitive tropical maize lines (22). A host of evidence suggests that maize cultivation in northern latitudes was enabled by the selection of allelic variants that led to a reduction in photoperiod sensitivity to allow flowering under longer photoperiods (23; 24; 25; 26; 27; 28; 22). Some of the early flowering alleles that conferred an adaptive advantage in highland environments are the result of *mexicana* introgressions into highland maize (24). Maize first entered into the US via the Mexican highlands (20), and these early flowering alleles show further evidence of selection in northern latitudes (24), consistent with a likely role for *mexicana* introgression(s) in maize adaptation to shorter daylength. When introduced into Northern Europe, photoperiod-insensitive maize from the Northern US and Canada thrived as it was already pre-adapted to northern latitudes and lower temperatures (29). The genetic, physiological, and phenotypic basis of these adaptations, however, is quite limited.

Plant phospholipids, as well as other glycerolipids such as sulfolipids, galactolipids, and non-polar lipids such as triacylglycerols, are involved in plant responses to low temperatures. Phospholipid levels are increased in plants exposed to low temperatures (30) and levels of unsaturated fatty acids in glycerolipids are reduced (31; 32), which may help maintain the fluidity of cell membranes. Under stressful conditions, the proportions of differently shaped lipids are modulated to maintain membrane flexibility while preventing membrane leakage. For instance, phosphatidylcholines (PCs) are rectangular polar lipids that are well suited for the formation of bilayer membranes because the size of their glycerol backbone, choline headgroup and fatty acid tails are similar. By contrast, lyso-phosphatidylcholine (LPC) is a triangular PC with a single acyl group that cannot form a bilayer because its headgroup is much larger than its fatty acid (33). LPCs do allow for some membrane movement, but high LPC concentrations act as a detergent (34) and can facilitate cell leakage and damage at low temperatures, effects that would be prevented by higher PC levels. In cold-adapted maize temperate lines and *Tripsacum* species (a distant maize relative), genes involved in phospholipid biosynthesis show accelerated rates of protein sequence evolution, further supporting an important role for phospholipid metabolism across several species during cold adaptation (35). Finally, multiple phospholipids can bind to Arabidopsis (*Arabidopsis thaliana*) FLOWERING LOCUS T (FT) and accelerate flowering. Recent work has shown that phosphatidylglycerol binds and sequesters FT in companion cells in low temperatures, while higher temperatures release it to the shoot apical meristem (36). There, it interacts with certain species of PC, the most abundant phospholipid in plant cells (37), through an unknown mechanism (38). Consistent with this observation, glycerolipid levels in maize have predictive power for flowering time (39).

Here, we identified an adaptive teosinte *mexicana* introgression that alters highland maize phospholipid metabolism and leads to early flowering. Using genome scans and linkage mapping, we identified *High PhosphatidylCholine1* (*HPC1*), a gene encoding a phospholipase A1, as a driver of high PC levels in highland maize. Data from thousands of genotyped landrace testcrosses grown in common garden experiments at different elevations in México showed a strong genotype-by-environment effect at the *HPC1* locus, where the highland allele leads to higher fitness in the highlands and reduced fitness at lower elevations. Furthermore, we determined that the highland *HPC1* allele, which was introgressed from teosinte *mexicana*, was carried northward and is now present in maize cultivars grown in the Northern US and European Flint lines. These results suggest that the *HPC1* highland allele has a beneficial effect in cold, high-latitude environments, where early flowering is advantageous.

## Results

### Highland Mesoamerican maize shows high PC/LPC ratios and selection of highly unsaturated PCs

As lipids play an important role in adaptation to adverse environments, we quantified the glycerolipid levels of 120 highland and lowland landraces^1^ from Mesoamerica and South America (Fig. 1A-B, Sup. File 1). This diversity panel, hereafter referred to as the HiLo diversity panel, was grown in highland (2650 meters above sea level [masl]) and lowland (20 masl) common garden experiments in Mexico. We determined that Mesoamerican highland landraces have high PC/LPC ratios, particularly when grown in the highlands (Fig. 1A-B).

**Figure 1.**
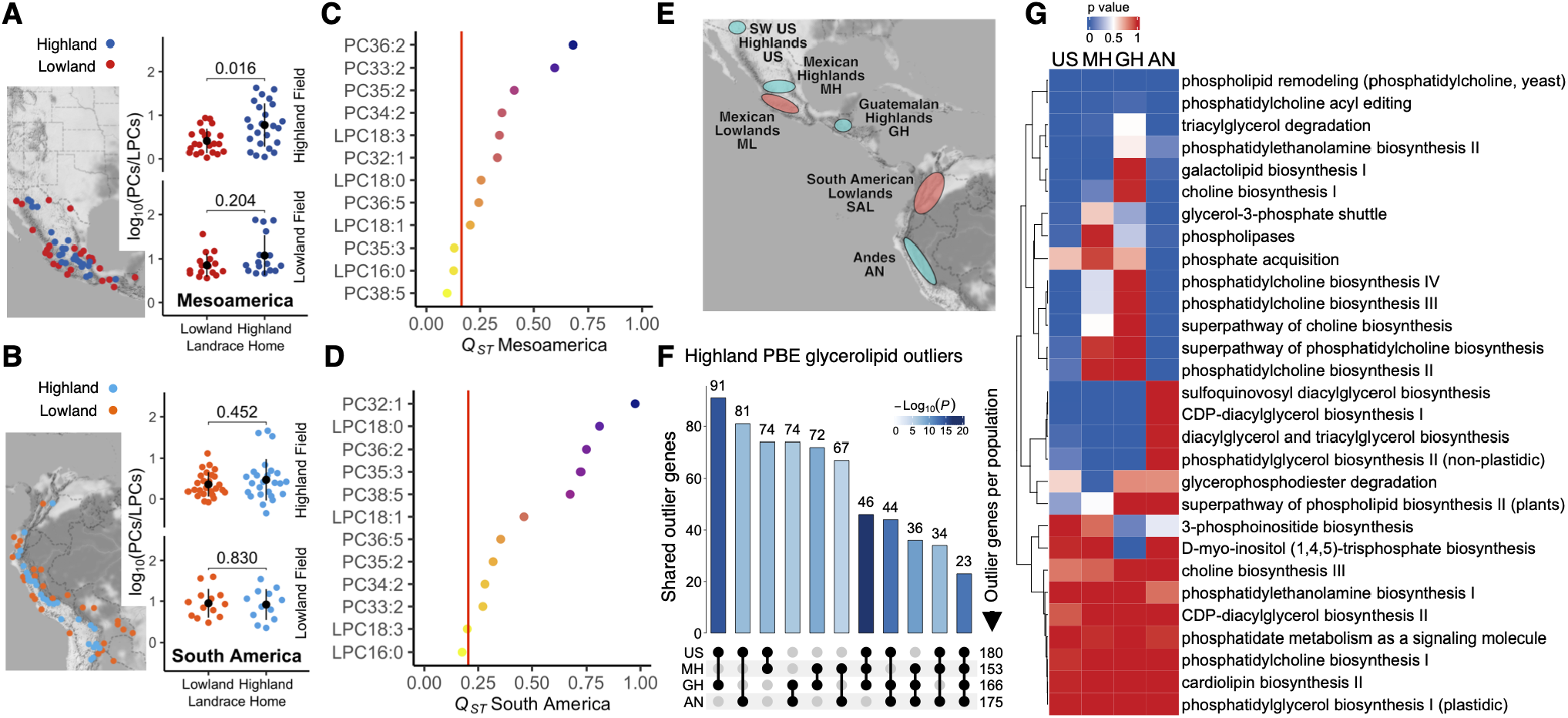
Phospholipid selection in highland maize. **A-B)** *Left:* Geographical origin of 120 maize accessions from the HiLo diversity panel used in the common garden experiment for glycerolipid quantitation. *Right:* PC/LPC ratio, log10 scaled, for highland and lowland landraces from (A) Mesoamerica, *n* = 54; and (B) South America, *n* = 53. The black cirlce indicates the mean, the vertical line is the standard deviation (SD). Significant differences were tested with a false-discovery rate (FDR) adjusted *t*-test; the resulting *p*-values are shown. **(C-D)** *Q_ST_-F_ST_* analysis of phospholipids between highland and lowland landraces from (C) Mesoamerica and (D) South America. Red line, neutral *F_ST_* + 2 SD. **(E-G)** Highland vs lowland Population Branch Excess (PBE) analysis. **(E)** PBE geographical sampling. **(F)** PBE outlier counts for glycerolipid genes in four highland populations. Bar shade indicates Fisher’s exact significance test for excess shared outliers. **(G)** Highland selection of glycerolipid-related pathways using PBE. The heatmap shows the probability of randomly sampling gene SNPs with mean PBE greater than the mean for the gene SNPs in each pathway.

The differences observed in phospholipid levels between highland and lowland maize may be the result of adaptive natural selection or random genetic drift during maize colonization of highland environments. To distinguish between these two possibilities, we compared the variance of each phenotype across the population with the genetic variance of neutral markers using a *Q_ST_* – *F_ST_* comparison (40). We calculated *Q_ST_* –*F_ST_* using diversity array technology sequencing (DartSeq) genotypes (41) from the same plants phenotyped for glycerolipid levels and calculated the *Q_ST_* –*F_ST_* values for each glycerolipid species for highland/lowland populations from each continent. Mean *Q_ST_* was greater than mean *F_ST_* in both Mesoamerican and South American comparisons. We identified PC and LPC species with higher *Q_ST_* values than the neutral *F_ST_* in both continents (Fig. 1C-D). The species with the highest *Q_ST_* value included long chain PCs with more than one unsaturation such as PC-36:2.

### Genes involved in PC/LPC conversion show strong highland selection signals

We selected a set of 597 maize glycerolipid genes from their functional annotations (see Materials and Methods) to identify selection signals using the Population Branch Excess (PBE) (42) statistic across four highland populations: Southwestern US (SWUS), Mexican highlands (MH), Guatemalan highlands (GH) and Andes (AN) (Fig. 1E) (19). We identified a significant excess of genes that are targets of selection in more than two populations (*p* < 3 × 10^−5^, 1F). The most over-represented intersection of selected glycerolipid genes was between the SWUS, MH and GH populations (*p* = 1 × 10^−15^, Fig. 1F), suggesting that genes were specifically selected in these three populations relative to the AN population and/or that there was closer kinship among SWUS, MH, and GH populations than the AN population and thus weaker statistical independence. From all annotated glycerolipid genes, 23 were consistently PBE outliers across all four highland populations (*p* =< 1 ×^−10^, Fig. 1F). We then assigned a PBE value to each of the 30 glycerolipid metabolism pathways (using a 10 kb window around each constituent gene) and compared their average PBE value with a genome-wide random sampling distribution of PBE values within genic regions. From these, we established that ‘phospholipid remodeling’ and ‘PC acyl editing’ exhibit significantly higher PBE values in all four populations, indicating a possible role for phospholipid remodeling in maize highland adaptation (Fig. 1G).

We considered two possible explanations for the extent of convergent selection in highland populations (see explanations in (19; 43)). Adaptation may be conferred by a small number of genes, thereby imposing a *physiological* constraint on the sources of adaptation leading to convergence. Alternatively, adaptation may be the result of many genes, but deleterious *pleiotropic* effects restrict the number of genes that can be targeted by selection, also leading to convergence. Using Yeaman’s *C_hyper_* statistic (43), which quantifies these two modes of convergent adaptation, we determined that the overlap among putative adaptive genes in the four highland populations cannot be explained merely by physiological constraints (*C_hyper_* = 3.96, Supplementary Table 1). A certain degree of pleiotropic constraint is therefore likely. Overlap between adaptation candidates was higher for the SWUS, MH and GH population pairs (*C_hyper_* = 4.79) than between the Andean and SWUS, MH and GH pairs (*C_hyper_* = 3.14).

To further understand selection at the gene level, we used genotyping by sequencing (GBS) data from 2,700 Mexican maize landraces, generated by the SeeD project (44; 15), to run a *pcadapt* (45) analysis to determine how loci might contribute to observed patterns of differentiation along major principal components of genetic variation. The first *pcadapt* principal component separated Mexican landraces based on the elevation of their geographical origin (Fig. 2B). Using this first principal component, we identified outlier single nucleotide polymorphisms (SNPs) across the genome that are significantly associated with genetic variation along elevation and potentially involved in local adaptation (Fig. 2A). From the list of 600 maize glycerolipid-related genes, 85 contained SNPs that were *pcadapt* outliers for association with the first genetic principal component (top 5% *log*_10_(*P*)), and of which eight were also PBE outliers for Mexican highlands (Fig. 2A, Supplementary table 2, Sup. File 2). These eight selection candidates, supported by two different sources of evidence, included two genes coding for putative enzymes whose orthologs are known to directly catalyze PC/LPC interconversion reactions. The first gene, Zm00001d039542, with a *log*_10_(*P*) of 110.28, encoded a putative phospholipase A1 that we name *High PhosphatidylCholine 1*, *HPC1*. The second gene, Zm00001d017584, with a *log*_10_(*P*) of 99.31, encoded a predicted *Lyso-Phosphatidylcholine Acyl Transferase 1* that we will refer as *ZmLPCAT1*. (Fig. 2C). Although these two types of enzymes catalyze broadly opposite reactions (degradation vs biosynthesis of PC) they are unlikely to catalyze strictly reverse reactions in the Lands cycle. A1 phopholipases attack the phosphatidylcholine at the *sn-1* carbon, while LPC acyl transferases usually acylate *sn-2* (46; 47). Instead, plant PLA1 enzymes like *HPC1* are better known for their role in the first step of jasmonic acid biosynthesis (48; 49).

**Figure 2.**
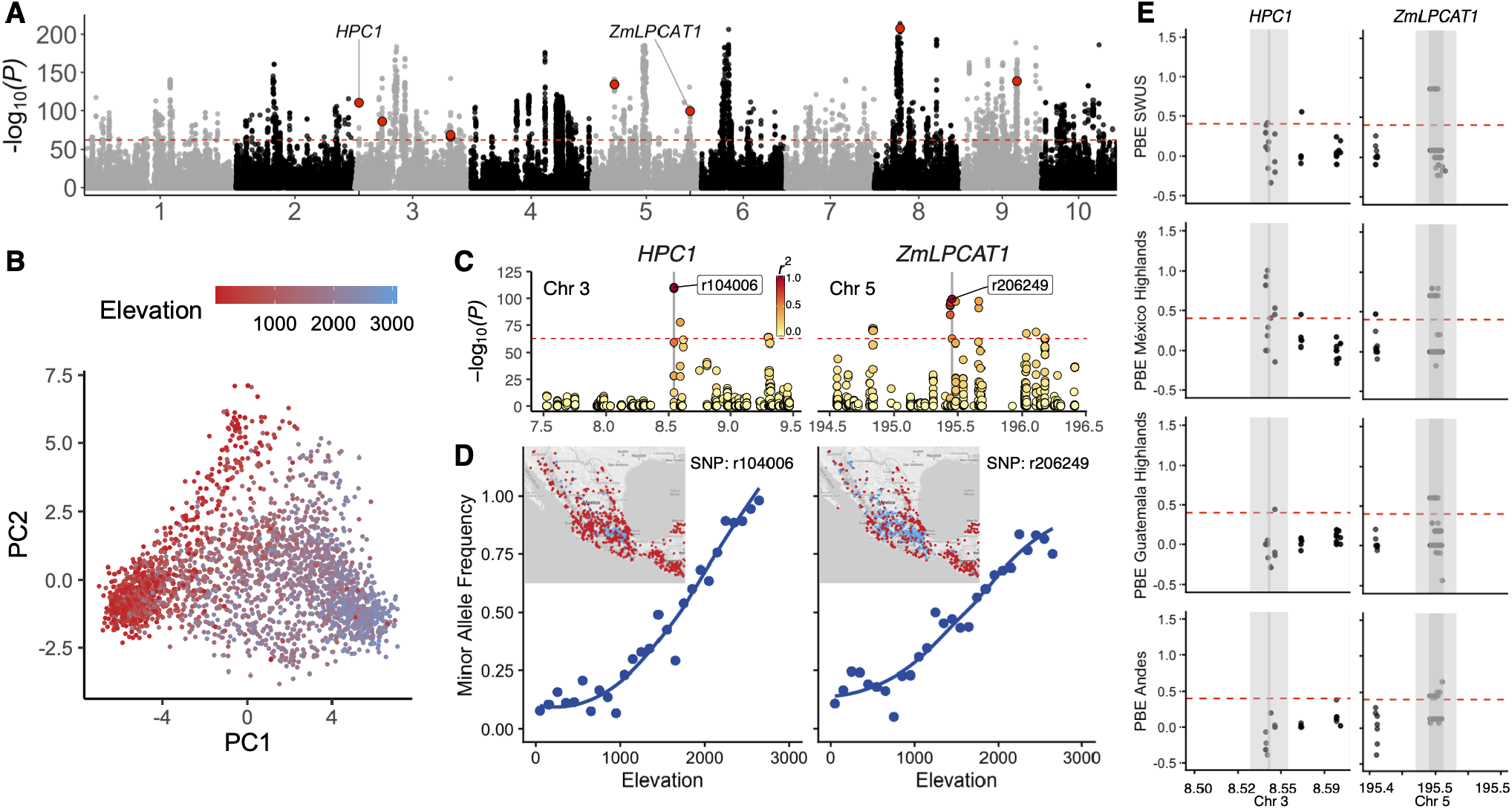
Evidence of highland selection in genes determining PC/LPC ratios. **(A)** Association with genetic principal component 1 (PC1) from *pcadapt* analysis of Mexican landraces (open pollinated varieties), dashed marks the top 5% *log*_10_(*P*). Red points show SNPs in glycerolipid metabolism genes that are also PBE outliers for Mexican highlands (Supplementary Table 2). From these, *HPC1* and *ZmLPCAT1* are the top genes with orthologs catalyzing PC/LPC interconversion. **(B)** Scatter plot of the *pcadapt* first two genetic principal components illustrating that PC1 correlates with elevation. **(C)** Extended region from (A) of the 1 Mb interval around *HPC1* and *ZmLPCAT1*. Linkage disequilibrium (*r*2) with the peak SNP for each gene are illustrated by the color scale; both peak SNPs are located in the coding sequence. **(D)** Elevation clines for the peak SNPs from (C), the insets show the geographic distribution of the highland (blue) and lowland (red) alleles. **(F)** Population Branch Excess of SNPs in *ZmLPCAT1* and *HPC1*. Dark grey, coding sequence; light grey, 10 kb upstream and downstream; dashed line is the threshold for the top 5% PBE value outliers.

Both genes showed strong changes in elevation-dependent allele frequency (Fig. 2D) across Mexican landraces. *HPC1* was not an outlier for branch excess between the Andean and the South American lowland accessions. By contrast, *ZmLPCAT1* was indeed a PBE outlier for all four populations, which may indicate parallel/convergent selection for this gene between Mesoamerican and Andean landraces. Importantly, both *HPC1* and *ZmLPCAT1* are annotated as part of pathways with outlier PBE values in all highland populations for ‘phospholipid remodelling’ and ‘PC acyl editing’ (Fig.Fig. 1F, Sup. File 2). Taken together, these two independent population genetic approaches show that pathways involved in phospholipid remodelling, including genes controlling the PC/LPC ratio like *HPC1* and *ZmLPCAT1*, show strong selection signals in highland maize. These results indicate that selection on phospholipid metabolite levels (Fig. 1B-D) is supported at the gene-level by outlier PBE and principal component analysis values *pcadapt* in genes controlling phospholipid biosynthesis and degradation.

### A major QTL explaining PC to LPC conversion overlaps HPC1

To further characterize the genetic architecture of phospholipid biosynthesis in highland maize, we developed a recombinant inbred line (RIL) BC1S5 population derived from a cross between the temperate inbred line B73 and the Mexican highland landrace Palomero Toluqueño (PT), using B73 as the recurrent parent (75% B73, 25% PT) (50).

The parental PT accession is a popcorn landrace (*palomero* means popcorn in Spanish) from the Toluca Valley in México (*Mexi5* CIMMYTMA 2233) (Fig. 3A). We grew the HiLo diversity panel and the B73 x PT BC1S5 RIL population in the same highland and lowland common gardens and collected samples for glycerolipid analysis. The locally adapted PT landrace displayed higher fitness than B73 in the highland field (Fig. 3A), probably due to adaptation to low temperatures in this highland environment. In the Mexican highlands, values of 5 GDDs per day are typical, while 15-20 GDDs/day are common in lowland environments.

**Figure 3.**
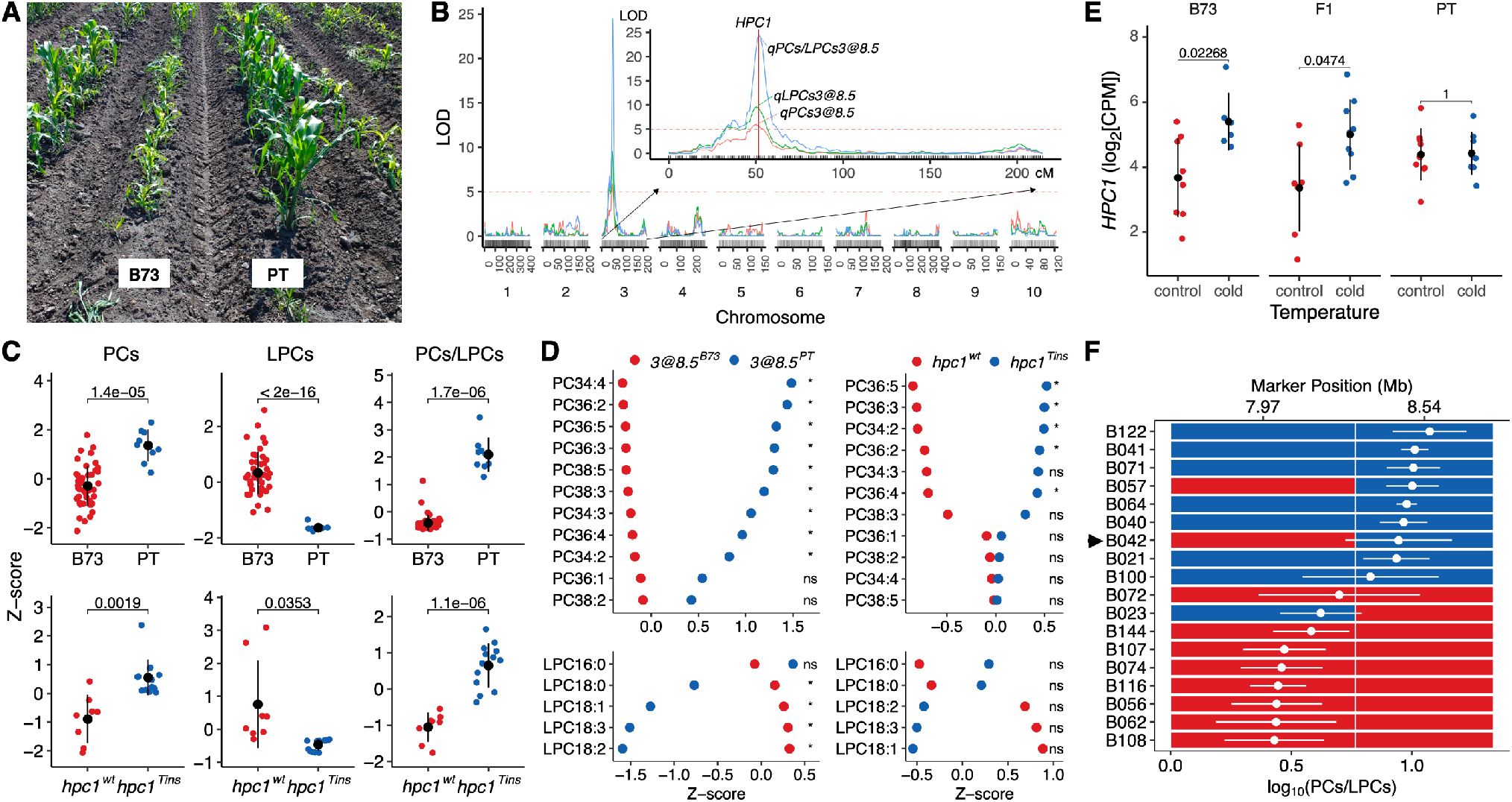
*HPC1* defines a major QTL explaining PC/LPC conversion. **(A)** PT and B73 plants growing in the highland Metepec field. (B) QTL analysis identified overlapping major QTLs around 8.5 Mb on chromosome 3 for PC and LPC levels and PC/LPC ratio, using data collected from plants grown in highland and lowland fields. The QTL peaks coincide with the physical location of *HPC1*. (C) Effect sizes for PCs, LPCs and PC/LPCs (z-score normalized) in RILs that are either homozygous for B73 or PT at 8.5 Mb on chromosome 3 (top row) and CRISPR-Cas9 *hpc1Tins* mutant and wild-type plants (bottom row). Significant differences were tested by *t*-test; the resulting *p*-values are shown. **(D)** Effect sizes for individual PC and LPC species (z-score normalized) in RILs at 8.5 Mb on chromosome 3 (left) or the CRISPR/Cas9 *hpc1Tins* mutant (right). * Significant difference at *p* < 0.05 (*t*-test, after Benjamini & Hochberg correction), *ns* not significant. **(E)** *HPC1* expression analysis in B73, PT and their F_1_ hybrids grown in standard and cold temperatures in a growth chamber. Significant differences were tested by *t*-test with Benjamini & Hochberg correction; the resulting *p*-values are shown. **(F)** PC/LPC ratio for several RILs. RIL B042 (indicated by the black arrow) bears a recombination event 500 bp upstream of the *HPC1* translation start codon. In panels (C-F), phenotypes associated with the B73 haplotype are in red; the equivalent values for the PT haplotype are in blue.

We detected major quantitative trait loci (QTLs) for the sum of LPC species levels, PC species levels, and the PC/LPC ratio that all mapped to the same locus on chromosome 3, around 8.5 Mb (Fig. 3B). We tested for epistatic interactions for LPC levels, PC levels, and the PC/LPC ratio through a combination of R/qtl `scantwo` and `stepwise` functions (53). The three QTLs *qLPCs3@8.5*, *qPCs3@8.5* and *qPCs/LPCs3@8.5* were robust against environmental effects and were detected in both the highland and lowland environments. The additive effect of the PT allele at these QTLs was associated with high levels of PCs, low levels of LPCs, and consequently high PC/LPC ratios, while the B73 allele had the opposite effect (Fig. 3C, top panel). Individual PC and LPC species QTLs at this locus Fig. S2 showed the same additive effect for the PT allele as the sum of each class (PCs, LPCs, and PCs/LPCs) of species (Fig. 3C, top panel. All individual LPC QTLs at the *qLPCs3@8.5* locus corresponded to LPCs that contained at least one double bond in their fatty acid (Fig. S2, Sup. file 3). *qPCs3@8.5* was driven mainly by PC species with more than two fatty acid double bonds, such as PC 36:5 (Fig. 3D and Fig. S2 A-D, Supplementary file 3). We then sought to identify candidate genes underlying the QTLs on chromosome 3. The PC/LPC ratio QTL had the highest significance, with a logarithm of the odds (LOD) of 24.5, and explained the most phenotypic variance (87%). The underlying QTL interval contained 72 genes within its 1.5 LOD drop confidence interval (7.9-10 Mb). We hypothesized that the metabolic phenotypes we observed might be due to a gene involved in PC-LPC conversion. The maize genome encodes 75 putative phospholipases (Fig. S3A), of which half are predicted to be phospholipase A1-type (PLA1) (Fig. S3B). Notably, *HPC1* mapped within the interval (position on chromosome 3: 8,542,107 bp to 8,544,078 bp), making it a high confidence candidate causal gene (Fig. 3B). HPC1 was predicted to have phospholipase A1-Igamma1 activity and can be classified as a PC-hydrolyzing PLA1 Class I phospholipase based on its two closest Arabidopsis orthologs (encoded by *At1g06800* and *At2g30550*) (54). PLA1-type phospholipases hydrolyze phospholipids in the *sn-1* position and produce a lyso-phospholipid and a free fatty acid (Fig. S3B). In B73, *HPC1* was one of the most highly expressed phospholipase genes, with expression almost exclusively restricted to vegetative leaves (V4-V9) (Fig. S4A) (55), which was the biological material we sampled for glycerolipid analysis. In B73 leaves, *HPC1* was also the most highly expressed gene within the QTL interval (Fig. S3C) (55). Class I phospholipases are chloroplast-localized proteins; in agreement, we identified a chloroplast transit peptide (CTP) at the beginning of the predicted HPC1 sequence using the online tool ChloroP (56). We validated the chloroplast localization of HPC1 by transiently expressing a construct encoding the HPC1 CTP fragment fused to green fluorescent protein (GFP) in *Nicotiana benthamiana* leaves (Fig. S5).

The effect of *HPC1* on PC/LPC levels may be caused by misregulation of *HPC1* expression in highland landraces and/or by a mutation affecting HPC1 enzymatic activity. To distinguish between these two possibilities, we analyzed *HPC1* expression in B73, PT, and the corresponding F_1_ hybrid plants grown at high and low temperatures to simulate highland and lowland conditions, respectively (Fig. 3E). Under cold conditions, *HPC1-B73* was up-regulated, but *HPC1-PT* was not (Fig. 3E). The lack of up-regulation in cold conditions of *HPC1* may explain the high PC/LPC levels in PT. However, *HPC1* expressed to the same levels in B73 and PT under control conditions. In the F_1_ hybrids, *HPC1* expression was consistent with a dominant B73 effect. We also observed a dominant B73 effect at the metabolic level when we analyzed B73 x PT RILs that are heterozygous at the *HPC1* locus (Fig. S3D). Variation in *HPC1* may also affect enzymatic activity of the HPC1-PT variant. To test this hypothesis we sequenced three B73 x PT RILs (B021, B042, B122) that are homozygous for the *HPC1-PT* allele. We discovered a recombination point between 493 and 136 bp upstream of the *HPC1* translation start codon (Fig. 3F, Fig. S6) in RIL B042, resulting in a chimeric locus with the coding region from PT combined with a promoter segment from B73. PC/LPC levels in RIL B042 were similar to other RILs that are homozygous for the PT haplotype at the 8.54 Mb marker at the QTL peak (Fig. 3F). This result supports the hypothesis that the metabolic effect in the B73 x PT RILs is likely due to an impaired function of the HPC1-PT enzyme rather than to changes in the *HPC1-PT* regulatory region. However, regulatory variants in the first 500 bp of the promoter may also impact expression levels in RILB042.

If *HPC1* is the underlying causal gene of this QTL, the observed metabolic phenotypes would be consistent with a reduction or loss of HPC1-PT enzyme function, leading to higher levels of PCs and lower levels of LPCs in the PT background. To test this hypothesis we generated mutants in *HPC1* via clustered regularly interspaced short palindromic repeats (CRISPR)/CRISPR-associated nuclease (Cas9)-mediated genome editing (Fig. S7A) in the B104 inbred, a temperate stiff stalk inbred derived from the Iowa Stiff Stalk Synthetic population like B73. We identified two transgenic mutants, hereafter designated *hpc1^CR T ins^* and *hpc1^CR T del^* (Fig. S7A). We then measured PC and LPC species in wild-type and mutant plants grown under long day conditions. The phospholipid profiles of the *hpc1^CR^* plants replicated those of the *PT* allele in the RILs (Fig. 3 C-D bottom panels and Fig. S7B), confirming that the *HPC1-PT* allele impairs HPC1 function and thus underlies the QTL on chromosome 3 around 8.5 Mb. Lastly, we *in vitro* translated HPC1-B73 and HPC1-PT versions of the protein in a cell-free system and incubated them with various phospholipid substrates. We then measured the amount of phospholipid substrate and lyso-phospholipid product for each compound (Fig. S8). This experiment confirmed that both HPC1 variants have PLA1 activity and suggested that HPC1-B73 may have higher activity on substrates like PC36:4 that show stark differences in abundance between highland and lowland lines, as well as in *hpc1CR* lines. Together, the RIL and CRISPR mutant results showed that *HPC1* underlies a major metabolic QTL explaining PC/LPC ratio. A mutation in the flap-lid domain of HPC1 affecting substrate accessibility likely leads to impaired function in the highland PT allele.

### Identification of the putative causal SNP in HPC1

We Sanger sequenced the *HPC1* locus from several RILs harboring the PT haplotype at *HPC1* and identified several non-synonymous SNPs within the coding sequence that might influence HPC1 function (Fig. S11).

We focused our attention on SNP 631 affecting the flap lid domain, which led to a conservative replacement of valine by isoleucine (V211I, Fig. 4A). The flap lid domain is important for phospholipase activity and is located in a lipase class 3 domain (PFAM domain PF01764) that is highly conserved across the tree of life (52). We recovered 982 observations of the lipase class 3 PFAM domain from 719 prokaryotic species using PfamScan (62; 63) and estimated optimal growth temperatures from their tRNA sequences (64). We then tested whether genetic variation in the sequence encoding the lipase class 3 domain was significantly associated with optimal growth temperature in bacteria (51). We detected several significant associations, all of which were located in the flap lid region (Fig. 4A, bold letters). Notably, the presence of a valine at residue 211, as observed in the PT allele, was accompanied by lower optimal growth temperatures relative to an isoleucine at residue 211, as observed in B73 (Fig. 4B), suggesting that the PT allele may be better adapted to the low temperatures to which highland maize is exposed.

**Figure 4.**
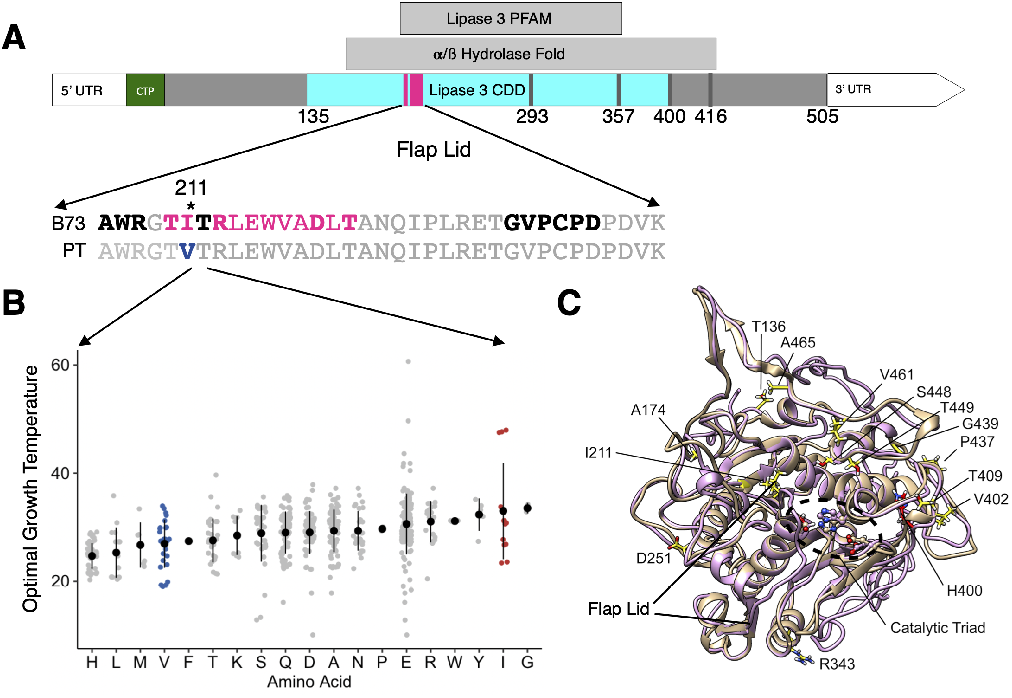
Analysis of the possible causal SNP affecting flaplid domain of HPC1. **(A)** *HPC1* coding sequence showing different encoded features. CTP, chloroplast transit peptide. The coding sequence fragment that encodes the flap lid domain and the corresponding amino acids are shown in magenta. Residues in bold showed significant associations with prokaryote optimal growth temperature (51; 52). **(B)** Optimal prokaryote growth temperature (OGT) test of different amino acids at the residue equivalent to the polymorphic residue 211 in HPC1. **(C)** Top I-TASSER model (purple) for the putative phospholipase encoded by *HPC1-B73*, overlaid on chain B of the crystal structure of phospholipase A1 from Arabidopsis (PDB 2YIJ, tan). Residues that differ between the PT and B73 phospholipase are shown in yellow and some are labeled.

We used the crystal structure of phospholipase A1 from Arabidopsis (PDB 2YIJ) to model the structure of HPC1-B73 by I-TASSER (Fig. 4C). Residues that differ between HPC1-PT and HPC1-B73 are shown in yellow. The catalytic triad and H400 identified from the conserved domain database (CDD) conservation analysis are also labeled. Our models placed H416 rather than H400 in the catalytic triad. Of the residues that differ between HPC1-PT and HPC1-B73, residues 211, 448, and 449 were the closest to the catalytic triad. I211 was positioned on the N-terminus of the flap lid domain and is well suited to stabilize binding of a lipid substrate through hydrophobic interactions. Replacement of I211 with V211 results in the loss of a methyl group that may influence the strength of these hydrophobic inter-actions, affect substrate binding, and/or the dynamics of the flap lid domain. These results strongly point to the mutation in the flap lid domain as the most likely underlying mutation affecting HPC1 activity.

### HPC1 shows strong elevation-dependent antagonistic pleiotropy in Mexican landrace fitness phenotypes

Our selection and QTL analysis provided strong evidence that *HPC1* is under selection in highland maize and controls phospholipid metabolism. To evaluate the possible fitness effects of HPC1 variation in locally adapted landraces across México, we re-analyzed phenotypic data from a previously reported F1 association mapping panel (44; 15) composed of about 2,000 landrace F1s grown in 23 common garden environments across an elevation gradient. We then fitted a model to estimate the effect of variation at *HPC1-PT* on the relationship between fitness traits and elevation (57). *HPC1* was a clear outlier in a genome wide association study (GWAS) of genotype-by-elevation fitness traits like flowering time and yield (Fig. S9A-B), indicating that elevation-dependent variation at *HPC1* not only has an effect on phospholipid levels but also on fitness traits. Indeed, variation at *HPC1* showed significant genotype elevation effects for several fitness traits (Fig. 5A). The effect of *HPC1* on flowering time revealed antagonistic pleiotropy between highland and lowland environments (Fig. 5A). The highland *HPC1-PT* allele was associated in low elevations with delayed flowering, increasing days to anthesis (DTA) by about one day. Meanwhile the same allele exhibited accelerated flowering at high elevation (with a decrease in DTA of almost one day, Fig. 5A). Variation at *HPC1* also displayed conditional neutrality on fresh ear weight and grain weight per hectare traits: the highland allele had no effect in lowland environments but was associated with greater values in highland environments (Fig. 5A). We also checked previous reports for associations between *HPC1* and flowering time in other populations through the MaizeGDB (61) genome browser. We in fact found a significant flowering time SNP in the *HPC1* coding sequence (Fig. 5B) for the Nested Association Mapping (NAM) population (60). This additive flowering time locus is only 6 bp from the focal SNP we used to test *G E* at *HPC1* in the SEEDs panel (Fig. 4). Variation at this SNP correlated with a reduction in flowering time of 8.5 hours, relative to B73, and explained 1.12% of the trait variance, which is about one third of the largest effect observed for flowering time variation in the NAM population. We then used genetic marker data from the HapMap 3, which includes the NAM parents, (66) to analyze linkage disequilibrium (LD) of the *HPC1* region (Fig. 4B). We detected a strong LD block of about 150 bp in length in the coding sequence that includes the focal SNP mentioned above (Fig. 5A,B). We identified another LD block covering the 5’ region of *HPC1* and the promoter region up to 2 kb upstream of *HPC1* (Fig. 5B). Interestingly, this second LD block on the promoter overlapped with two strong ATAC-seq (assay for transposase-accessible chromatin followed by sequencing) peaks identified in B73 ((59), Fig. 5B). These results confirmed that the SNPs associated with fitness traits like flowering time on the HPC1 coding sequence are not linked to other SNPs upstream of the HPC1 coding sequence. However, the SEEDs data set lacks GBS markers for several kb upstream of *HPC1*, raising the possibility of a second regulatory variant in the promoter (Fig. 4B) that might have an effect on *HPC1* expression. We further evaluated the possible effect of *HPC1* on flowering using both *hpc1^CR^* mutants in long-day conditions during the Summer of 2021 in Raleigh, NC. Although the mutants showed high PC/LPC ratios, we observed no significant difference in flowering time relative to the wild type (Fig. S7C). As shown in (Fig. 4A) the effect of the *HPC1* allele on flowering time has a strong G x E pattern and we only observe a significant effect in very high or low elevations. We speculate that the absence of significant differences in the B104 CRISPR mutants in Raleigh conditions could partly be explained by the strong G X E effect of *HPC1* and/or genetic background effects.

**Figure 5.**
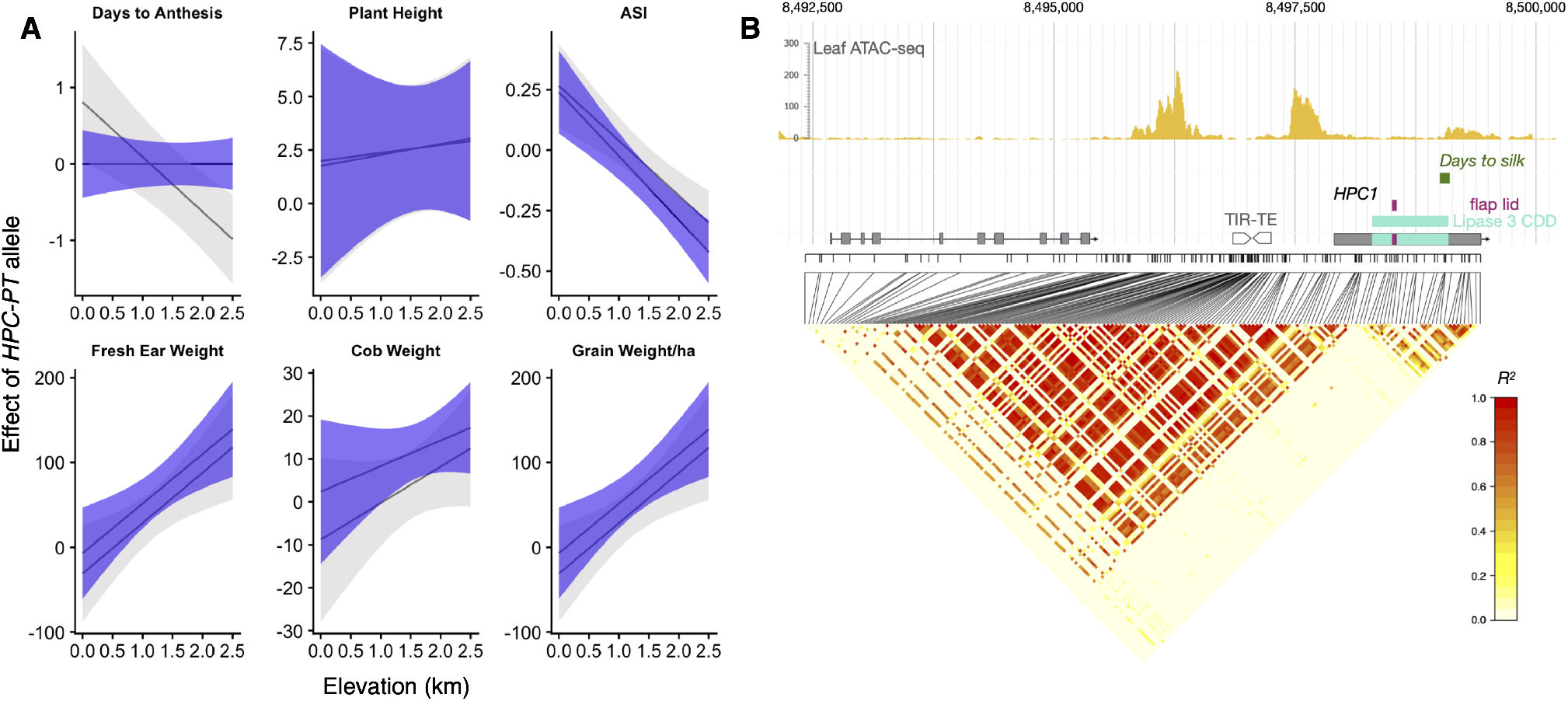
Fitness effects of the *HPC1-PT* allele and *HPC1* LD analysis. **(A)** We used best linear unbiased predictions (BLUPs) and GBS data from 2,700 landrace topcrossess from (15), evaluated in 23 common gardens at different elevations in México. We modeled each trait as a function of the *HPC1-PT* genotype, trial elevation, and tester line, with controls for main effects and responses to elevation of the genomic background. Gray lines and ribbons show estimates of the effect of the highland allele of *HPC1-PT* as a function of common garden elevation ± 2 standard errors of the mean, using the *GridLMM* package (57). Purple lines show estimates of the *HPC1-PT* effect in a model that also adjusts for effects of days to anthesis. ASI, anthesis to silking interval. **(B)** Linkage disequilibrium and genomic features of *HPC1* and upstream region. LD heatmap drawn with (58) from HapMap 3 data. Top track shows leaf ATAC-seq peaks in B73, data from (59). The regions coding for the lipase and flap-lid domains are highlighted on the HPC1 gene model. See 4A for further details. Days to Anthesis indicates the GWAS SNP on the lipase domain from (60). TIR-TE, terminal inverted repeat transposable element. Tracks obtained from MaizeGDB B73 V5 browser (61).

In summary, the SNPs in the short, lipase-domain-encoding LD block of *HPC1* show strong genotype x elevation fitness effects in both Mexican landraces grown across multiple altitudes and the NAM population. The phospholipid changes induced by *HPC1* have physiological effects that may explain the strong selection of *HPC1* in highland environments.

### HPC1-PT was introgressed from teosinte mexicana and is conserved in Flint inbred lines

We explored the segregation of the V211I SNP among other highland maize varieties. We detected the PT allele at high frequencies in highland landraces from México and Guatemala. In addition, the PT allele segregated in Southwestern US landraces. The B73 allele was fixed in lowland Mexican, South American, and Andean landraces (Fig. 6A). These results were consistent with our PBE results (Fig. 2F). The PT allele was also present in one fourth of all teosinte *parviglumis* accessions tested and in both *mexicana* accessions reported in Hapmap 3 (66) (Fig. 6A). This observation prompted us to examine whether the PT allele was the result of post-domestication introgression from teosinte *mexicana* during maize highland colonization, or whether it was selected from *parviglumis* standing variation. To test for introgression from *mexicana*, we used *f_d_* data from (10) and established that the genomic region containing *HPC1* shows signatures of introgression from *mexicana* into highland maize (Fig. 6B). We then performed a haplotype network analysis using SNP data from the *HPC1* coding region of 1,160 Mexican accessions from the SeeD Dataset (44) that are homozygous for all SNPS across the coding region and the teosinte inbred lines (TILs) from Hapmap 3 (66). We identified nine haplotype groups that cluster mainly based on elevation (Fig. 6C). The two major groups, II and VI, contained mainly lowland and highland landraces, respectively. The two *mexicana* teosinte inbred lines (TIL08 and 25) belonged to group IV (Fig. 6C) together with highland landraces primarily collected in the Trans-Mexican Volcanic Belt (30/36 from the highlands of Jalisco, Michoacán, México, Puebla, and Veracruz). We then examined whether this *mexicana* haplotype (denoted *ZxHPC1*) that is introgressed into Mesoamerican highland maize was also present in modern maize inbred lines.

**Figure 6.**
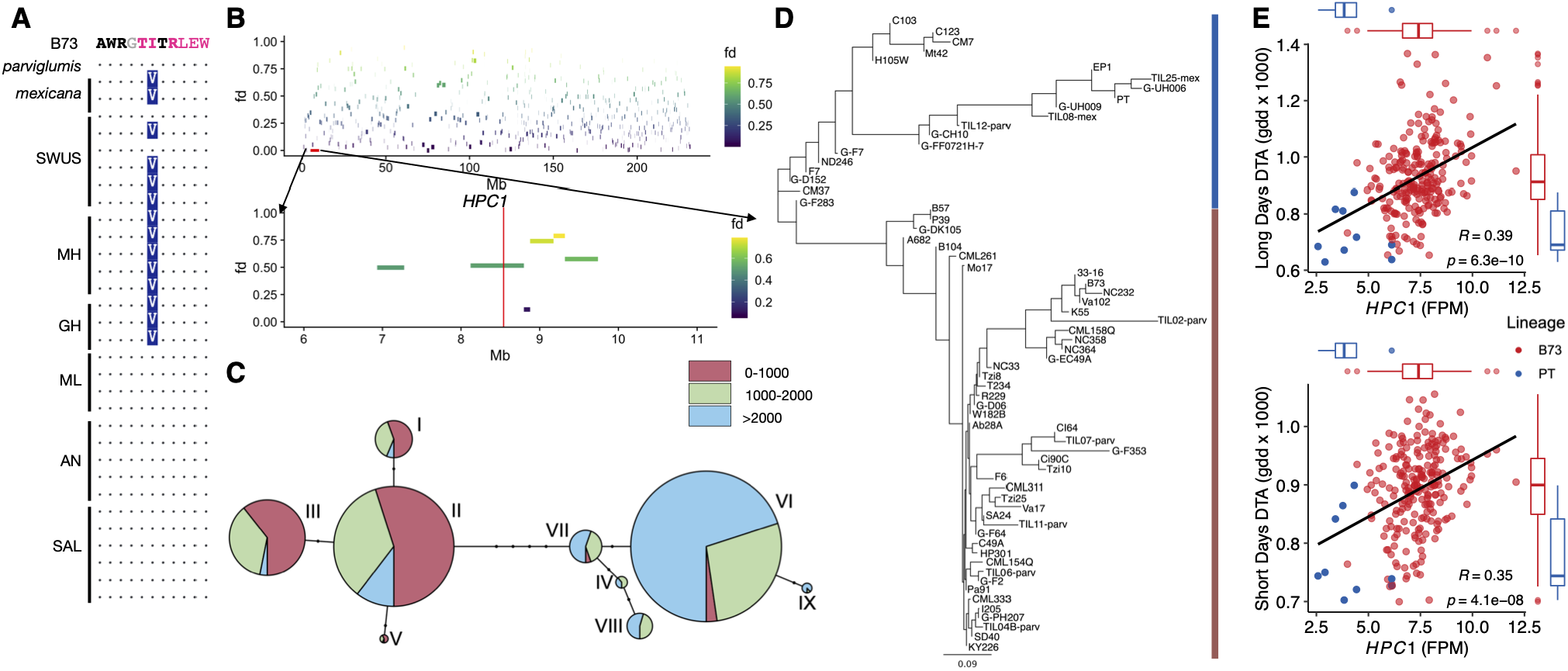
Introgression of teosinte *mexicana* into maize *HPC1*. **A)** Alignments around the V211I polymorphism in the flap-lid domain of HPC1 in B73, *mexicana* and *parviglumis* and landraces of the Southwestern US (SWUS), Mexican Highland (MH) and Lowlands (ML), Guatemalan Highlands (GH), Andes (AN) and South America Lowlands (SAL). **B)** *f_d_* analysis of the *mexicana* introgression. Data were obtained from (10). **C)** Haplotype network analysis of SNPs within the *HPC1* coding region, using 1,060 Mexican homozygous individuals from the SeeD dataset. Haplotypes are color-coded by elevation: red: 0-1000 masl, green: 1000-2000 masl, blue >2000 masl. **D)** Cluster analysis of the *HPC1* coding region using a sample of Hapmap3 inbred lines and the PT landrace. **E)** Correlation between *HPC1-PT* expression and days to anthesis (DTA) in plants grown in short or long days. Inbred lines from the PT lineage shown in panel C are indicated in blue and inbred lines from the B73 lineage are in red; data from (65).

To this end, we performed a neighbor-joining cluster analysis using Hapmap 3 (66) inbred lines including those from the 282 inbred panel, Teosinte inbred lines, German lines, and PT. We identified two main groups, one containing the *HPC1-PT* haplotype and the other containing the *HPC1-B73* haplotype. PT and the teosinte *mexicana* inbred lines TIL08 and TIL25 clustered together with Northern European Flint inbred lines such as EP1, UH008, and UH009 (Fig. 6D). Other Northern US flints, e.g., CM7, were also closely related to the *mexicana ZxHPC1* haplotype. These data suggest that after introgression into highland maize, the *ZxHPC1* haplotype was maintained in Flint materials adapted to cold environments in the Northern US, Canada, and Europe. We then used a large gene expression dataset consisting of multiple developmental stages of the 282-maize diversity panel (65) and phenotypic datasets collected from the same panel grown in long days and short days. Notably, *HPC1* expression levels were highly correlated with flowering time in both long and short days. Lines carrying the *HPC1-PT* allele were characterized by lower *HPC1* expression and earlier flowering times relative to lines carrying the *HPC1-B73* allele (Fig. 6E). Taken together, these data show that *HPC1* was introgressed from teosinte *mexicana* into highland maize and that this introgression was carried over into high latitude-adapted modern inbred lines with low *HPC1* expression and early flowering.

### HPC1/Phosphatidylcholine interactions with maize flowering time protein ZCN8

Using the same expression dataset from the 282 panel (65), we discovered that *HPC1* and *ZmLPCAT1* expression levels are inversely correlated in most tissues (Fig. S12), further supporting the idea that these two enzymes are coordinately regulated. We observed significant associations between *HPC1* expression levels in aerial tissues and several flowering time traits (Fig. S12) The magnitude of these associations was similar to that seen for other well-characterized flowering time genes (Fig. S12) such as *ZEA CENTRORADIALIS8* (*ZCN8*) and the APETALA2(AP2)/ETHYLENE RESPONSE FACTOR (ERF) transcription factor gene RELATED TO AP2.7 (*ZmRAP2.7*). In Arabidopsis, florigen (FT) has recently been shown to interact with both PC and phosphatidylglycerol (PG) depending on the ambient temperature and the cellular location of FT (38; 36). As ZCN8 is a homolog of FT, it may mediate the hastening of flowering time seen in highland maize. Recent work improved upon the crystal structure of Arabidopsis FT, onto which we modeled ZCN8 and compared the PC binding sites (Fig. S13A,B) (67). Using AutoDock Vina to simulate docking and the sites from (67), we identified sites 2 and 4 and 1 and 4 as potential binding sites of PC34:2 and PC36:2 with site 4 heavily favored in both cases (Fig. S13A-C). As maize ZCN8 and Arabidopsis FT are highly conserved, the predicted binding site similarity is not surprising (Fig. S13D). We next heterologously produced and purified ZCN8 fused to a SPOT tag from yeast (*Saccharomyces cerevisiae*) cells to assess lipid binding to ZCN8 *in vivo*. We extracted and analyzed all lipids co-purifying with the recombinant protein following the same lipidomics pipeline used for maize lipids. We identified PC34:2 in the purified ZCN8 protein in all samples confirming experimentally that ZCN8 effectively bound PC species (Fig. S14).

## Discussion

Understanding the genetic, molecular, and physiological basis of crop adaptation to different environments and the role that wild relatives have played in these processes is relevant for the identification of favorable genetic variation that can be used to improve modern crops. The repeated events of maize adaptation to highland environments constitute an excellent natural experiment to study local adaptation. Recent studies (19; 18; 14) have helped expand our understanding of the genetic basis underlying maize highland adaptation. However, the responsible molecular, physiological, and genetic mechanisms underlying maize highland adaptation and the possible role of highland maize traits in modern, commercial varieties remain largely unknown. Phospholipids are key structural components of plant membranes that also function as signaling molecules during adaptation to stresses that would be prevalent in highland environments (54; 68) such as low phosphorus availability (69; 70; 71) and low temperatures (30; 31; 72). In Arabidopsis and rice (*Oryza sativa*, phospholipid species regulate flowering time via interactions with *Arabidopsis* FT and rice Heading date 3a (Hd3a), respectively (38; 36; 73). Flowering time is a major driver of maize adaptation to highland environments (44; 15; 74) and to northern latitudes (22; 21).

Genes involved in the biosynthesis and degradation of phospholipids appeared to have been repeatedly selected in several highland maize populations of North America, Central America, and South America (Figures Fig. 1 and Fig. 2). *ZmLPCAT1* and *HPC1* were two such genes with the strongest, repeated signals of selection, as measured by PBE and *pcadapt* in highland populations (Fig. 2). In a previous study, *ZmLPCAT1* exhibited high *F_ST_* values when highland and lowland landraces were compared (18). In temperate inbred lines, *HPC1* was up-regulated while *ZmLPCAT1* was down-regulated by cold stress (Fig. S4B-C) (75). These expression patterns are in agreement with the high and low PC/LPC ratios observed in our experiments. Furthermore, we determined here that *HPC1* is up-regulated in B73 and B73 x PT F_1_ hybrids after cold exposure. By contrast, the *HPC1-PT* and *HPC1-B73* alleles were expressed to comparable levels in control conditions, and the *HPC1-PT* allele was not induced upon cold conditions (Fig. 3E). Selection at these two loci is likely to have driven the high PC/LPC ratio we observed in highland Mexican landraces (Fig. 1).

Our QTL analysis of the PC/LPC ratios in a B73 x PT mapping population and in the *hpc1^CR^* mutant alleles demonstrates that the highland *HPC1-PT* allele results in an enzyme with impaired function that alters highland Mexican maize PC metabolism, leading to higher PC/LPC ratios (Fig. 3, Fig. S7B). Adaptive loss-of-function mutations can be an effective way to gain new metabolic functions in new environmental conditions (76). Taking advantage of the conserved lipase domain of HPC1 in bacteria, we used a new method that can identify how genetic variation in DNA regions encoding protein domains is correlated with optimal growth temperature of bacteria (51; 52). The probable causal SNP in HPC1 changed an amino acid in the flap-lid domain of HPC1 (Fig. 5A) that may affect substrate accessibility and/or substrate binding (Fig. 5A). Indeed, the flap-lid domain has been the target of biotechnological modification for lipases (77). The region encoding the flap-lid domain was located at a recombination point that separates two clear LD blocks. The first LD block covered a 2-kb promoter region of *HPC1* and the first 600 bp of *HPC1*, while the second covered the rest of the *HPC1* coding sequence. In fact, this pattern is characteristic of selective sweeps that leave two LD blocks on either side of the adaptive mutation sweep (78). LD analysis together with the open chromatin detected by ATAC-seq that overlaps the same promoter region (Fig. 4B) and the correlation between *HPC1* expression and flowering time (Fig. 6E) presented here suggest a possible transcriptional regulation of *HPC1* that may act additively with the coding sequence variation we described.

Why were the metabolic changes induced by HPC1 variation selected for in highland maize? PC metabolism is intimately connected to multiple stress responses and developmental pathways; alterations in PC amounts and PC/LPC ratios affect overall plant fitness. The *qPC/LPC3@8.5* QTL is driven by individual QTLs for PC and LPC species with high levels of unsaturated fatty acids (Fig. 3D). Several of these species, like PC 36:5 and LPC 18:1 (Fig. 3D, Sup. File 3), have been shown to display similar patterns during Arabidopsis cold acclimation (31) and sorghum (*Sorghum bicolor*) low temperature responses (72). PC 36:5 also showed high *Q_ST_* values when comparing highland and lowland landraces from both Mesoamerica and South America (Fig. 1C-D, Sup. File 5). In maize, *HPC1* expression is under the control of the circadian clock (79) with a peak at the end of the day. In Arabidopsis, highly unsaturated PC (34:3, 34:4, 36:5, 36:6) species increase in the dark (80). This peak in contents coincides in maize with low *HPC1* expression levels during the same dark hours (79). PC, and lipid metabolism in general, is also intimately connected to flowering time. For instance, PCs were shown to bind to Arabidopsis FT in the shoot apical meristem to hasten flowering (38; 67) by unknown cellular mechanisms. Similarly, PG species can sequester FT in phloem companion cells in low temperatures (36) and then release FT into the phloem later after temperatures increase, allowing FT reach the shoot apical meristem. In agreement with an effect of lipids on flowering time, overexpression of a gene encoding a secretory phospholipase D delayed heading time in rice (73). In line with a role for phospholipids in flowering, we established that genetic variation at SNPs within the region of *HPC1* encoding the lipase domain exhibits a strong interaction with elevation for the highland *HPC1-PT* allele. This variation leads to a delay in flowering time in low elevations and an acceleration at high elevations, both of which close to one day in amplitude. Interestingly, the effect of the highland *HPC1* allele exhibited typical conditional neutrality in yield-related traits, with higher fitness conferred by the *HPC1-PT* allele in highlands (Fig. 4A). In the NAM population, we identified another SNP mapping to the region encoding the lipase domain that is associated with a hastening of flowering time by eight hours with respect to B73 (60), further supporting the role of *HPC1* in controlling flowering time. Analysis of *HPC1* CRISPR mutant alleles in the B104 background grown in Raleigh displayed a PC/LPC phenotype that mimicked the highland allele (Fig. S7A-B) but not a flowering time phenotype, probably due to the strong G x E effect of *HPC1* and/or genetic background effects.

The strong G x E effect we observed in *HPC1* is similar to the well-known teosinte *mexicana* introgression *inv4m* (14). In fact, our analysis showed that *HPC1* is indeed another introgression from teosinte mexicana (Fig. 6A-C). Recent analysis using sympatric teosinte and maize populations across elevation gradients in Mexico further supports the introgression of *mexicana* at *HPC1* and shows that the *mexicana* ancestry of *HPC1* increases at a rate of +0.079 per 100 m of elevation (81). We further demonstrated that the mexicana intogression at *HPC1* is conserved in high latitude-adapted Flint lines from both Europe and the USA (Fig. 6D). *HPC1* in inbred lines carrying the highland *ZxHPC1 mexicana* haplotype was expressed at low levels and resulted in earlier flowering (Fig. 6E) (65).

Adaptation to higher latitudes involved a reduction of photoperiod sensitivity and flowering time that enabled maize to thrive in longer day conditions characteristic of the growing season at high latitudes (22; 21; 27; 26). Additive mutations in the regulatory region of the gene *ZCN8* (82), including a teosinte *mexicana* introgression, lead to higher expression of *ZCN8*, which contributes to maize adaptation to long days in temperate conditions (83). *ZCN8* is a close ortholog of Arabidopsis *FT*, whose encoding protein interacts with several species of phospholipids to modulate flowering time (38; 36). A similar interaction was also demonstrated for the rice FT ortholog Hd3a, (73) and we hypothesize that the same may be occurring in maize. Comparison of docking simulations of phospholipids with ZCN8 using the Arabidopsis FT crystal structures as a model (67) showed similar PC interactions in ZCN8 (Fig. S13). We corroborated this interaction via mass spectrometry analysis of lipids bound to ZCN8 heterologously produced in yeast (Fig. S14).

In summary, we used a combination of genomic scans, linkage mapping, lipidomics and reverse genetics to identify and clone the adaptive gene *HPC1*, introgressed from teosinte *mexicana*, in highland maize landraces. HPC1 variants lead to a major reorganization of phosphatidylcholine metabolism. We showed that the fitness advantage conferred by the *HPC1* highland *mexicana* allele is due, at least in part, to its association with flowering time. This effect may have contributed to adaptation of maize to colder, higher latitudes.

Our work is the first to identify the important role of a gene controlling phospholipid metabolism in plant local adaptation and further supports the emerging role of phospholipid metabolism in fine-tuning flowering time across different plant species (38; 36; 83). This study highlights the largely underappreciated role of highland maize and highland teosinte *mexicana* in modern maize.

## Materials and Methods

The diversity panels and mapping populations used in this manuscript for population genetics measures of selection, QTL mapping and G x E analysis have been described previously by (19), (15), (44), (41) and (50). Lipidomics analysis were performed with high resolution mass spectrometry UPLC-MS. Mutant alleles of HPC1 were obtained using CRISPR-CAS9 editing. Full description and details of all materials and methods is provided on the SI Appendix. All the data and code is contained within the paper, supplementary material and associated Github repository of the project.

## Acknowledgments

This work was supported by Conacyt Young Investigator (CB-238101), Conacyt National Problems (APN-2983), UC-Mexus and NC State startup funds awarded to Rubén Rellán-Álvarez. Fieldwork and mapping population development was supported by NSF-PGR award 1546719 to Jeffrey Ross-Ibarra, Ruairidh Sawers, Daniel Runcie and Matthew Hufford. Allison Barnes was supported by NSF-PGRP PRFB grant 2010703. Andi Kur was supported as an AgBioFEWS fellow by the NSF award 1828820. Fausto Rodríguez-Zapata were supported by the Science and Technologies for Phosphorus Sustainability (STEPS) Center, a National Science Foundation Science and Technology Center (CBET-2019435). This study made use of NMRbox: National Center for Biomolecular NMR Data Processing and Analysis. A Biomedical Technology Research Resource (BTRR), which is supported by NIH grant P41GM111135 (NIGMS). This work was performed in part by the Molecular Education, Technology and Research Innovation Center (METRIC) at NC State University, which is supported by the State of North Carolina. We thank Josh Strable, Peter Balint-Kurti, Luis Herrera-Estrella and James Holland for critical review of the manuscript and feedback. We thank the members of ther Rellán-Álvarez, Sawers, Hufford and Flint-Garcia labs for their help developing and evaluating mapping populations. We thank Cruz Robledo and the Puerto Vallarta Agricultura Invernal team, including members of the local Wixárika community for their work in our lowland field. We thank Fernando Delgado, Denise Costich and Cristian Gálvez and the staff of the CIMMyT Metepec field station and the CIMMyT Germplasm bank for their assistance in genetic nurseries and field evaluation. We thank the and CIMMYT and Puerto Vallarta Winter Nursery crews that have helped generate and evaluate the mapping populations used in this manuscript. We specially want to acknowledge the indigenous people of the Americas and the ingenuity through which they domesticated and made the spread and adaptation of maize throughout the continent possible. The field experiments performed in this work in México and North Carolina were performed on land that originally belonged to Native Americans. This work would not have been possible without the international maize research community and the willingness of so many colleagues to support the development of new research programs.

## Supplementary Information

### Supporting Information Text

#### Materials and Methods

##### A. Populations used in the analysis

Highland and lowland populations used for Population Branch Excess analysis consisted of three to six accessions from each of the highland and lowland populations and have been previously described in (1, 2). The 120 landraces from the HiLo diversity panel were selected and ordered from the CIMMYT germplasm bank to maximize a good latitudinal gradient sampling across Mesoamerica and South America. For each highland landrace (>2,000 meters above sea level [masl]) a lowland landrace (<1,000 masl) was selected at the same latitude (<0.5) to form 60 highland/lowland pairs, with 30 from each continent. The list of the accessions used is provided in Sup. file 1. B73 x Palomero Toluqueño (PT) recombinant inbred lines (RILs) were developed by crossing B73 with a single PT plant (Mexi5 accession, CIMMYTMA 2233) that was then backcrossed with B73 once and selfed five times (BC1S5). We used individual landrace accession genotype and fitness data from the CIMMYT Seeds of Discovery project (SeeD) (3) to calculate *pcadapt* (4) values and GxE effects of *HPC1-PT*.

##### B. Field Experimental Conditions and sampling

Two replicates of the HiLo diversity panel accessions and three replicates of the B73 x PT RILs were planted in a highland and lowland common gardens. The highland common garden was located in Metepec, Edo de México, (1913’28.7”N 9932’51.6”W) in the Trans-Mexican volcanic belt. The field is at 2,610 masl, and the range of average monthly temperatures along the year vary from 5C to 21.5C. The lowland common garden was located in Valle de Banderas, Nayarit, (2047’01.2”N 10514’47.0”W) in the Pacific Coast. The field is at 50 masl, and the range of average monthly temperatures along the year vary from 20C to 29C. Between growth stages V4 and V6, we used a leaf puncher to collect 50 mg of fresh tissue (10 discs) from the tip of the second leaf above the last leaf with a fully developed collar. Tissue discs were immediately flash-frozen in liquid nitrogen. We collected all samples from a field in a single day between 10:00 am and 12:00 pm, approximately 3 h after sunrise. Samples were transported in dry ice to the lab and stored at –80C until extraction.

##### C. Glycerolipid analysis

We crushed frozen samples in a Retsch tissue grinder (Haan, Germany) for 40 sec at a frequency of 30 pulses per second. We performed lipid extraction following Matyash and collaborators (5). First, we added 225 L of cold methanol (MeOH) to each sample. For the blanks, we prepared MeOH containing a Quality Control (QC) mix (Sup. File 6). We vortexed each sample for 10 sec, keeping the rest of the material on ice. Then, we added 750 L of cold methyl tert-butyl ether (MTBE). For the blanks, we used MTBE containing 22:1 cholesterol ester as internal standard (Sup. File 6). We vortexed each sample for 10 sec, followed by 6 min of shaking at 4C in the orbital mixer. We next added 188 L of LC/MS grade water at room temperature (RT), and vortexed samples for 20 sec. We centrifuged the samples for 2 min at 14,000 rcf and recovered 700 L of supernatant from the upper organic phase. We then split the supernatant into two aliquots of 350 L each, one aliquot was used for the generation of the lipid profile and the other aliquot was used to prepare pools that were used along the lipid profiling. Finally, we dried the samples using a speed vacuum concentration system. We resuspended dried samples in 110 L of MeOH-Toluene 90:10 (with the internal standard CUDA, 50 ng/mL). We vortexed samples at low speed for 20 sec and then sonicated them at RT for 5 min. We then transferred aliquots of 50 L per sample into an insert within an amber glass vial. The UHPLC-QTOF MS/MS utilized were Agilent 1290 and Agilent 6530, respectively. We used a Waters Acquity charged surface hybrid (CSH) C18 2.1×100 mm 1.7 m column which we initially purged for 5 min. We coupled the UHPLC column with a VanGuard pre-column (Waters Acquity CSH C18 1.7m). We injected six “no sample injections” at the beginning of each run to condition the column, followed by ten samples, one pool (made out of the mix of the second aliquot of all the samples contained per UHPLC plate) and one blank. We injected 1.67 L per sample into UHPLC-QTOF MS/MS ESI (+); the running time per sample was 15 min. Mobile phase “A” consisted of 60:40 acetonitrile:water, 10 mM of ammonium formate and 0.1% formic acid. Mobile phase “B” consisted of 90:10 isopropanol:acetonitrile, 10 mM ammonium formate and 0.1% of formic acid. The flow rate was 0.6 mL/min and the column compartment was maintained at 65° C. Initial conditions were 15% B; the gradient uniformly increased until reaching 100%. At 12.10 min the mobile phase composition returned to initial conditions. We used the mass spectrometer (Q-TOF MS/MS) in positive electrospray ionization mode (ESI). The source parameters were: ESI gas temperature 325 C, nebulizer pressure 35 psig, gas flow 11L/min, capillary voltage 3500 V, nozzle voltage 1000 V, and MS TOF fragmentor and skimmer 120 and 65 V, respectively. We used a mass range between 60 and 1,700 m/z, under acquisition parameters. As for reference mass parameters, we used a detection window of 100 ppm and a minimum height of 1,000 counts. We performed a retention time (rt) correction of the acquired data using Agilent MassHunter Qualitative Analysis B.06.00 version and Microsoft Excel. To extract ion chromatograms (EICs) of the internal standards within the run we used Agilent MassHunter Qualitative Analysis. We identified the time of the highest intensity point of each EIC, which we used as the current retention time of the experiment. We used the method retention time for internal standards and the current rt and we fitted a polynomial regression to calculate new retention times using retention times from 501 lipids of a MS1 m/z-rt library (See Sup. File 7). In MSDIAL (6), identification of lipids is based on two approaches: the MSP file and MS/MS identification setting included in MSDIAL and the use of a post identification file containing accurate m/z and rt for a list of lipids. In this study we used both identification approaches. Under positive ion mode, the MSP file and MS/MS identification setting has a total of 51 lipid classes selectable for identification. The post identification file that we used was the retention time-corrected MS1-MS2 mz-rt lipid library that we explained before. We used MSDIAL (6) version 3.40. To use MSDIAL, we converted the raw data from .d to .abf format with Reifycs Abf converter (https://www.reifycs.com/AbfConverter/). Then, we filtered the MSDIAL alignment results based on whether compounds intensity was ten times above blank intensity. Next, filtered data were normalized using Systematic Error Removal using Random Forest (SERRF) (7). This normalization is based on the QC pool samples. We filtered out, now the normalized features, considering a coefficient of variation (CV) equal or less than 30% among the pools. To curate the data for duplicate features, isotopes and ion-adducts, we utilized MS-FLO (8). We also normalized the curated data using the sum of all known metabolite signal (mTIC). After data processing and normalization, we used lipid intensities for further analysis.

##### D. Glycerolipid analysis of *hpc1^CR^* mutants

Dry samples were resuspended in 110 L of 100% MeOH (with the internal standard CUDA, 50 ng/mL), vortexed at low speed for 20 s and then sonicated at RT for 5 min. We transferred the samples into amber glass vials with inserts prior to analysis. Lipid profiling was performed using a Thermo Scientific Orbitrap Exploris 480 mass spectrometer coupled to a Thermo Vanquish Horizon UHPLC. Chromatographic separation was achieved using the same guard and analytical column type, mobile phase composition and gradient conditions described for UHPLC-QTOF analyses. Avanti Splash Lipidomix Mass Spec Standard (Avanti Polar Lipids, Alabaster, Alabama) was used to check system suitability and a Lipid QC Standard Mixture (Sup. File 6) and pooled QC samples were used to monitor instrument performance. Methanol solvent blanks were injected at the beginning of the sequence and throughout the batch between approximately 10 injections and pooled QC samples were injected between every 6–8 sample injections. For ion source parameters the spray voltage was set at 3500 V, sheath gas 60 (arb), aux gas 15 (arb), sweep gas 2 (arb) and ion transfer tube and vaporizer temp at 350 C. Full scan and data-dependent MS/MS (ddMS^2^) scans were collected in positive ESI mode following 2 *μ*L sample injections. Full scan spectra were obtained from *m/z* 220–1700 using an Orbitrap resolution setting of 120,000, RF lens set at 25%, normalized AGC target at 25% and maximum injection time set to auto. ddMS^2^ scans were performed using a cycle time of 0.6 sec, intensity threshold of 5.0e4, and a 3 sec dynamic exclusion using a 10 ppm mass tolerance. Additional parameters for ddMS^2^ experiments were as follows: Isolation window equal to 1 Da, stepped collision energy at 20 and 30%, Orbitrap resolution set to 15,000, scan range set to auto, normalized AGC target of 100% and a maximum ion injection time of 55 msec. Data files were uploaded to LipidSearch 4.2.2 (Thermo Scientific, Tokyo, Japan) for identification of PC and LPC species. Skyline-daily (9) was then used to obtain normalized peak areas for the identified PCs and LPCs using the internal standards PC(12:0/13:0) and LPC(17:0), respectively.

##### E. Q_ST_ – F_ST_ analysis of glycerolipid data

To maximize independence between traits for *Q_ST_* analysis we clustered the phospholipid traits based on correlation values into 20 groups with complete-linkage hierarchical clustering. Single metabolites were then selected from each cluster, resulting in 20 minimally correlated traits figure:Sup:lipid*_c_lusters.Quantitativetraitdifferentiation*(*Q_ST_*) was contrasted to the distribution of *F_ST_* for neutral genetic markers (10). Genotypes were obtained previously from the same plants used for glycerolipid analysis (11). A Bayesian regression model (R package brms, (12)) was used to partition phenotypic variance between population pairs (Mesoamerican highland/Mesoamerican lowland, South American highland/South American lowland):

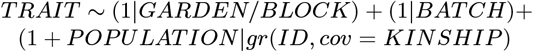

In this model, TRAIT value is modeled by random effects BLOCK within GARDEN, BATCH, and INDIVIDUAL (which share covariance modeled by the additive relationship matrix KINSHIP, produced by R function A.mat, R package sommer, (13)) within POPULATION. This model was run with adapt_delta = 0.99 (default 0.8), 4 chains, and 6000 iterations (default 4000). For 6 of the 40 models, 6000 iterations were insufficient to reach convergence, so this number was increased to 10000 (and in one case, to 14000). From these models, posterior samples were extracted (R function posterior_samples) and used to acquire the within-population and between-population variance components required to calculate *Q_ST_*:

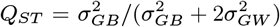

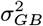 and 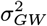 are the betweenand within-population genetic variance components, respectively (14). We report the median *Q_ST_* for each population contrast of each trait. Pairwise *F_ST_* was calculated with the R function fst.each.snp.hudson (R package *dartR*, 15).

##### F. *pcadapt* analysis of biological adaptation in Mexican landraces

In order to conduct genome scans for signatures of adaptation we used the *pcadapt* (4) package. *pcadapt* identifies adaptive loci by measuring how strongly loci are contributing to patterns of differentiation between major axes of genetic variation. Under simple models *pcadapt* captures major patterns of *F_ST_* but is conducted in a way that does not require population delimitation (16). As the genome scan comparison requires a focal SNP to be compared to the first K principal components of the genotype data, it can be biased by large regions of low recombination that drive the major axes of variation in the principal components. Thus, when SNPs from these low recombination regions are compared against principal components driven by linked loci spurious signals may arise. To prevent this bias from occurring, we used custom scripts to calculate the principal component step separately based upon all the chromosomes except for the chromosome of the focal SNPs being tested. The genotype data we used for this analysis was GBS data from roughly 2,000 landraces of Mexican origin collected by CIMMYT (www.cimmyt.org) as part of the SeeDs of discovery initiative (https://www.cimmyt.org/projects/seeds-of-discovery-seed/). From this, we calculated the strength of association between each SNP and the first five principal components (excluding the chromosome of the focal SNP) using the communality statistic as implemented in *pcadapt* version 3.0.4.

##### G. Glycerolipid pathways selection

We compiled a list of genes pertinent to glycerolipid metabolism starting with a search of all genes belonging to the *Zea mays* ‘Glycerophospholipid metabolism’ and ‘Glycerolipid metabolism’ KEGG pathways (17) (map identifiers: zma00564 and zma00561). With the NCBI Entrez gene identifiers in KEGG we retrieved the AGPv4 transcript identifiers used in Corncyc 8.0 (18, 19) from an id cross reference file found in MaizeGDB (18). This resulted in a list of 300 genes comprising 51 Corncyc pathways. Then we discarded Corncyc pathways tangentially connected to the KEGG glycerolipid metabolism list (sharing just one enzyme with the initial KEGG list) or that we judged to belong to different biological processes (e.g ’long chain fatty acid synthesis’, ’anthocyanin biosynthesis’). Finally, we added manually the ‘phosphatidylcholine biosynthesis V’ pathway that was missing. The list of 30 selected Corncyc pathways included genes outside the initial KEGG search results and raised the number of genes to 557. In addition to these, 37 genes were found to have an enzymatic activity related to phospholipid metabolism but not placed into any particular pathway, i.e orphan enzymes, consisting mostly of alcohol dehydrogenases. Sixteen additional genes found in KEGG were not annotated at all in Corncyc probably due to differences between AGPv4 and RefSeq pseudogene annotation of the maize genome. The list of all possible candidates coming either from KEGG or Corncyc that were orphan enzymes or were unannotated in Corncyc amounted to 597 genes (Sup. File 2). This process is documented in the `0_get_glycerolipid_genes.R` script of the `pgplipid` R package accompanying this paper (20).

##### H. Population Branch Excess Analysis

Population Branch Excess quantifies changes in allele frequencies in focal populations relative to two independent “outgroup” populations. We used *Zea mays spp. parviglumis* as one of the outgroup populations for all four highland groups. The other outgroup was Mexican lowlands in the case of Southwestern US, Mexican highlands and Guatemalan highlands; and South American lowlands in the case of the Andes population. We used calculated PBE SNP values for the four populations (described in detail in (1)) and we tested for selection outliers SNPs in the 594 phosphoglycerolipid-related candidate genes and the 30 Corncyc pathways (556 genes). We first defined PBE outlier SNPs as the top 5% of the PBE score distribution; this fraction corresponds to approximately 50000 out of 1 million genotyped SNPs in each population. Following (1), we defined a gene as a PBE outlier if it contained an outlier SNP within the coding sequence or 10 kb upstream/downstream. Then we tested for over-representation of genes selected in particular subsets of populations using Fisher’s exact test with the 32,283 protein-coding genes from the maize genome as background. For each pathway, we first selected all SNPs within coding regions and 10 kb upstream and downstream of genes in the pathway and we calculated the mean pathway PBE score. We then constructed a null distribution by drawing 10,000 samples without replacement of *n* SNPs from those found within 10 kb upstream and downstream of all protein-coding genes and we obtained the mean PBE for this null distribution. With the set of PBE outliers for glycerolipid metabolism in the four populations, we tested for evidence of physiological or pleiotropic constraint using the 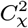 statistic (21).

##### I. QTL analysis of phospholipid levels

We analyzed glycerophospholipid QTLs in a mapping population of 57 RILs (BC1S5) from the cross B73 x PT. These RILs were grown in a highland site in Metepec, Edo de México at 2,600 masl during the summer of 2016 and in Puerto Vallarta, Jalisco at 50 masl during the winter of 2016/17. We analyzed the samples using UHPLC-QTOF, as above, and 67 leaf lipid species were identified. For QTL analysis we calculated the mean across all fields of individual lipid mass signal. We also used as phenotypes the sum total of the following lipid classes: diacylglycerols, triacylglycerols, PCs and LPCs. Furthermore, we also included the log base 10 transformed ratios of LPCs/PCs and the ratios of their individual species. We did a simple single marker analysis with “scanone“ using Haley-Knott regression, and assessed the QTL significance with 1,000 permutations.

##### J. CRISPR-Cas9 editing of and analysis of the effect of *hpc1^CR^* mutant on flowering time

CRISPR/Cas9 was used to create a mutant through Agrobacterium *Agrobacterium tumefaciens*-mediated transformation of background line B104 (22, 23). RNA guides (gRNAs) were designed as described in (24) for the B73 reference genome v4. The B104 and B73 sequence for were identical. The gRNA cassette was cloned into pGW-Cas9 using Gateway cloning. Two plants from the T0 transgenic events were identified through genomic PCR amplification and Sanger sequencing and were self-pollinated. In the next generation, several plants were allowed to self. Plants were genotyped using forward primer CAGTTCTCATCCCATGCACG and reverse primer CCTGATGAGAGCTGAGGTCC. Cas9 positivity was tested for using 0.05% glufosinate ammonium contained in Liberty herbicide or by PCR genotyping. *hpc1^CR^* T2 plants grown in Raleigh, NC under long day conditions were used for flowering time and lipid analysis. Wild-type plants from the same segregating families were used for analysis.

##### K. Cell-free protein production and substrate analysis

*HPC1-B73* and *HPC1-PT* coding sequences were synthesized without the fragment encoding the chloroplast transit peptide and codon optimized for *in vitro* translation in wheat. Protein was produced using the WEPRO7240 Kit from CellFree Sciences (Yokohama, Japan). Briefly, RNA was generated from the DNA templates using reverse transcription. Cooled RNA reactions were then mixed with creatine kiinase and WEPRO 7240 extract. This mixture was layered with translation buffer and incubated for 20h at 15°C. The bilayer was mixed and protein snap-frozen in liquid nitrogen before storage at −80 °C. Protein was incubated with various phospholipids for one hour in a buffer of 250mM Tris-HCl, 0.7M NaCl, and 10mM CaCl2, pH 7.4.

##### L. Infrared matrix-assisted laser desorption electrospray ionization mass spectrometry (IR-MALDESI-MS) analysis

LC-MS grade acetonitrile, water, and formic acid for the electrospray solvent and microscope slides were purchased from Fisher Scientific (Nazareth, PA, USA). The IR-MALDESI-MS platform has been described previously (25, 26). A 2970 nm laser (JGMA, Burlington, MA, USA) desorbed neutral molecules from the sample with a single burst of 10 pulses fired at 10 kHz (1.2 mJ/burst). The orthogonal electrospray plume consisted of 60:40 (v/v) acetonitrile/water with 0.2% formic acid. The electrospray flow rate was fixed at 2.0 μL/min and a 3600 kV potential was applied to the emitter tip. The IR-MALDESI source was coupled to an Orbitrap Exploris^TM^ 240 mass spectrometer (Thermo Fisher Scientific, Bremen, Germany) for high resolution, accurate mass analysis. The automatic gain control was disabled, and the ion accumulation time was fixed at 25 ms. The mass spectrometer collected mass spectra in positive mode over *m/z* 175 – 1000. The multi-injection RF threshold fixed at 6 to inject ions over the entire *m/z* range of interest through the S-lens for the duration of the ion accumulation time. The EASY-IC (fluoranthene, *m/z* 202.0777) was used for internal calibration to obtain parts per million mass measurement accuracy. The mass spectrometer resolution was fixed at 240,000_FWHM_ at *m/z* 200. The RF-level applied to the S-lens ion guide was fixed at 70% for all analyses. RastirX software paired with an Arduino microcontroller (Arduino, Ivrea, Italy) was used to coordinate electronic signals required for each analysis (27). For each IR-MALDESI-MS analysis, 1 μL of sample was pipetted onto a glass microscope slide and placed on the sample stage with the focal point of the laser focused at the top of the sample droplet. Mass spectra were viewed in QualBrowser directly. For data analysis and visualization, the raw datafiles were converted from the .RAW formal to .mzML files using MS Convert within the ProteoWizard software package (28). The .mzML files were then converted to .imzML files using imzMLConverter (29). The .imzML files were converted to .xlsx files using MSiReader v1.03c to effectively compile multiple analyses onto a single Microsoft Excel spreadsheet (30, 31).

##### M. Expression analysis of in B73, PT and B73xPT F1 hybrids in conditions simulating highland environments

Gene expression data were generated from leaf tissue from B73, PT and the B73xPT F1 hybrids. Plants were grown following the same protocol as in (32). Briefly, kernels were planted in growth chambers set to imitate spring temperature conditions in the Mexican lowlands (22°C night, 32°C day, 12 hr light) and highlands (11°C night, 22°C day, 12 hr light). Leaf tissue was sampled from the V3 leaf the day after the leaf collar became visible, between 2 and 4 hr after dawn. Tissue was immediately placed in a centrifuge tube, frozen in liquid nitrogen, and stored at –70°C. RNA-seq libraries were constructed, sequenced, and analyzed following (32). Briefly, randomly primed, strand specific, mRNA-seq libraries were constructed using the BRaD-seq (33) protocol. Multiplexed libraries were sequenced on 1 lane of an Illumina HiSeq X platform. Low quality reads and adapter sequences were removed using Trimmomatic v.0.36 (34), and the remaining paired reads were aligned to the maize reference genome B73 v4 and quantified using `kallisto` v.0.42.3 (35). Gene counts were normalized using the weighted trimmed mean of M-values (TMM) with the *calcNormFactors* function in *edgeR* (36) and converted to log2CPM.

##### N. Sanger sequencing of in RILs homozygous at the *qPC/LPC3@8.5* locus

We identified 3 RILs homozygous for the PT allele at the locus and we developed 6 sets of primers to sequence by Sanger across the coding region and the promoter. The location of primers along the locus are shown in Sup. file 8.

##### O. Association of with agronomic traits

We re-analyzed phenotypic data from the F1 Association Mapping (FOAM) panel of Romero-Navarro *et al* (37) and Gates *et al* (3) to more fully characterize association signatures of. Full descriptions of this experiment and data access are described in these references. We downloaded BLUPs for each trait and line from Germinate 3, and subset the data to only those lines with GBS genotype data from México. We fit a similar model to the GWAS model used by (3) to estimate the effect of the *HPC1-PT* allele on the traits’ intercept and slope on trial elevation, accounting for effects of tester ID in each field and genetic background and family effects on the trait intercept and slope using four independent random effects. We implemented this model in the *R* package *GridLMM* (38). We extracted effect sizes and covariances conditional on the REML variance component estimates and used these to calculate standard errors for the total *HPC1-PT* effect as a function of elevation. To test whether the phenotypic effects of *HPC1-PT* on yield components might be explained as indirect effects via flowering time, we additionally re-fit each model using days to anthesis (DTA) as a covariate with an independent effect in each trial.

##### P. Identification of features and domains of HPC1

The domains of were analyzed from UniProt identifier A0A1D6MIA3. The Lipase3–PFAM - PF01764, and alpha/beta Hydrolase fold were identified using InterPro, and Lipase3-CDD, shown in cyan, including flap lid, shown in magenta, and the S293, D357, and H400 catalytic triad were identified from CDD. H416 was identified as a substitute for H400 in the catalytic triad by protein modeling.

##### Q. Bacterial optimal growth temperature association with HPC1 flap-lid domain allelic variation

We compared the maize HPC1 protein sequence to prokaryotes with the same sequence to determine whether the identified residue change in maize and accompanying association with low temperature survival was consistent with observations in other organisms. We identified the Pfam domain PF01764 in the B73 protein sequence using the HMMER3 web server, we also identified 982 observations of the PF01764 Pfam domain in 719 prokaryote species using PfamScan (39, 40). We predicted the optimal growth temperature of these species using tRNA sequences as in (41). We aligned the maize and prokaryote PF01764 domain sequences with hmmalign from the hmmer3 package (42), and the aligned Pfam sequences were recoded to reflect nine amino acid physicochemical properties (43). We filtered the sequences to remove gaps in the domain alignment and observations with only partial domain sequences, then we clustered them based on sequence similarity, resulting in two clusters of observations within the domain. For each cluster, positions in the filtered alignment were associated with prokaryote optimal growth temperatures using a linear regression with all nine amino acid physicochemical properties. Seventeen sites in and around the flap-lid region of the protein passed a 10% FDR significance threshold, including the single residue change V211I previously identified in *HPC1-PT*. Welch’s two-sided t-test was used to compare the optimal growth temperatures of prokaryote species with the same (V) *HPC1-B73* aminoacid residue to the optimal growth temperatures of prokaryote species with the same (I) *HPC1-B73* allele at this site.

##### R. Subcellular localization of

We fused the HPC1-B73 chloroplast transit peptide (CTP, 52 aminocacids) to GFP. Three constructs encoding subcellular localization markers were used as control; cytoplasm (C-GFP), nucleus (N-GFP), and chloroplast (P-GFP). All markers were under the control of the cauliflower Mosaic Virus (CaMV) 35S promoter. These constructs were transiently infiltrated in *Nicotiana benthamiana* leaf cells.

##### S. Teosinte introgression in highland maize

To evaluate if the *HPC1-PT* allele is the result of standing variation from teosinte or an introgression from we used Patterson’s *D* (44) statistic and genome-wide *f_d_* (45) to calculate ABBA-BABA patterns. The data were obtained from (46). The material used in (46) included whole-genome sequence data from three highland outbred individuals: two PT and one Mushito de Michoacán; three lowland landraces: Nal Tel (RIMMA0703) and Zapalote Chico (RIMMA0733) obtained from (2) and BKN022 from (47); two inbred lines: TIL08 and TIL25; three inbred lines: TIL01, TIL05, TIL10 and *Tripsacum* TDD39103 (47) as an outlier.

##### T. Haplotype network analysis of in Mexican maize landraces and teosintes

We extracted SNP genotypes for from the TIL teosinte accessions in the HapMap 3 imputed data (47) and the 3700 Mexican landraces in the SeeD dataset. With the set of 1060 accessions that were homozygous at all sites in this genomic region we calculated a haplotype network depicting the minimal spanning tree for haplotypes covering 90% of the input accessions with the R package pegas (48), and haplotype frequencies for three elevation classes in the landraces (0–1,000; 1,000–2,000 and >3,000 masl).

##### U. Clustering analysis of in maize inbred lines and teosintes

Using v3 of the B73 genome, SNPs were obtained from PT (this paper), the 282 diversity panel (49), teosinte inbred lines and the German inbred lines from HapMap 3 (47). The selection was made to have a good representation of tropical, temperate and European inbred lines together with teosintes and PT lines. SNPs were aligned using Geneious2020.0.5 and a neighbor-joining cluster analysis was generated. To facilitate visualization and interpretation of the tree we condensed cluster branches from lines with identical haplotypes and from similar geographic locations. The full tree is available as Sup. file 9.

##### V. Expression analysis of candidate genes and association with flowering traits in the 282 diversity panel

We used gene expression RNA-seq data obtained from the 282 diversity panel at different developmental stages (50) and BLUP values of several flowering and photoperiod sensitivity traits (51) to study the correlation of expression values with flowering time traits.

##### W. Heterologous expression, purification of *ZCN8* using *Saccharomyces cerevisiae* and analysis of ZCN8-lipid binding

We synthesized *ZCN8* codon optimized for *Saccharomyces cerevisiae* and tagged with a SPOT peptide tag (Chromotek, Planegg, Germany); this construct will further be called ZCN8-SPOT. ZCN8-SPOT was then transformed into the galactose inducible vector pYES-DEST52 and expressed in *Saccharomyces cerevisiae* strain FY4. After 8 h of induction, cultures were harvested and frozen for future protein extractions. We purified protein with Chromotek SPOT-Cap beads (Chromotek, Planegg, Germany) and generally followed their protocol. Briefly, 50mL of induction pellet was resuspended in lysis buffer (10mM HEPES, pH 7.5, 150mM Nacl, 0.5mM EDTA pH 8.0, 1mM DTT, 10mM MgCl2, 1x HALT protease and phsophatase inhibitor (Thermo Fisher Scientific, Waltham, MA, USA), and 1% dodecylmaltoside) and ground in a Genogrinder with glass beads at max speed for one minute. This was followed by one minute on ice and repeated for a total of three minutes of grinding. The sample was then incubated on ice for five minutes, vortexed, and incubated for another five minutes. After pelleting, the supernatant was diluted with wash buffer (10mM HEPES, pH 7.5, 150mM NaCL, 0.5mM EDTA, pH 8.0, 1X HALT protease and phosphatase inhibitor, and 0.1% dodecylmaltoside) and put onto cleaned, equilibrated SPOT-Cap beads and left rotating at 4°C for 1 h. The beads and lysate were separated by centrifugation at 2500xg for 5 min and the beads were washed three times with 20x bead volume of wash buffer. Protein was eluted with either SPOT-peptide or 100mM glycine, pH 2.0. Glycine elutions were immediately neutralized with an equivalent amount of Tris base, pH 10.4. Elutions were frozen and lipids were extracted and analyzed at a later date as described above. The comparison data between an authentic standard and ZCN8-SPOT yeast was acquired with the same lipid profiling method as described in the Materials and Methods section for the Thermo Scientific Orbitrap Exploris 480 mass spectrometer with the following modifications: sample injection volume was increased to 10 *μ*L, full scan spectra were acquired from *m/z* 200 – 1000 and *m/z* 758.5694 was included in the target mass list for MS^2^ selection.

---

**Table S1.** Pairwise *C_hyper_* statistic for population comparisons. “population 1” and “population 2” are the two populations being compared.

| population 1 | population 2 | $C_{hyper}$ | $p$ |
| --- | --- | --- | --- |
| US | MH | 3.82 | 1.11E-04 |
| US | GH | 6.17 | 8.00E-10 |
| MH | GH | 4.37 | 1.24E-05 |
| US | AN | 3.51 | 3.37E-04 |
| MH | AN | 2.73 | 4.43E-03 |
| GH | AN | 3.16 | 1.16E-03 |

**Table S2.**
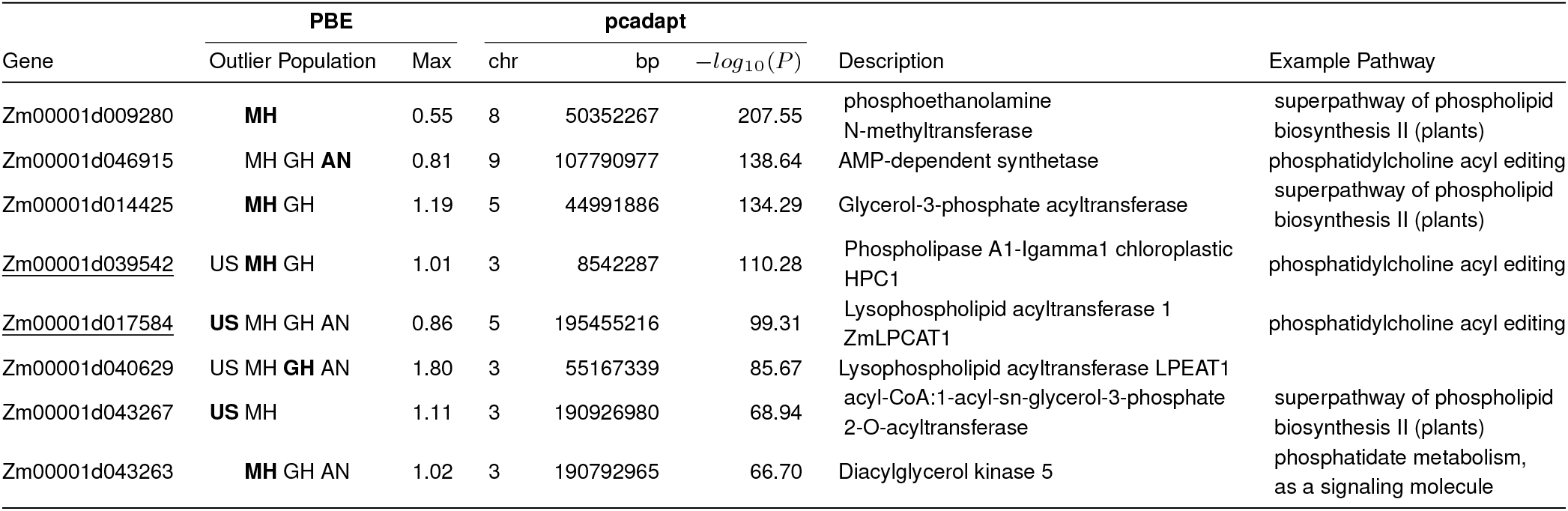
Selection outliers simultaneously detected in PBE and pcadapt analysis. These phosphoglycerolipid metabolism genes are in the top 5% of Mexican highlands PBE, and are also in the top 5% of significance (−*log*_10_(*P*)) for the association with pcadapt PC1 (elevation loaded principal component). Further populations where the gene is an outlier for PBE are listed as well, maximum PBE population is highlighted in bold.

**Fig. S1.**
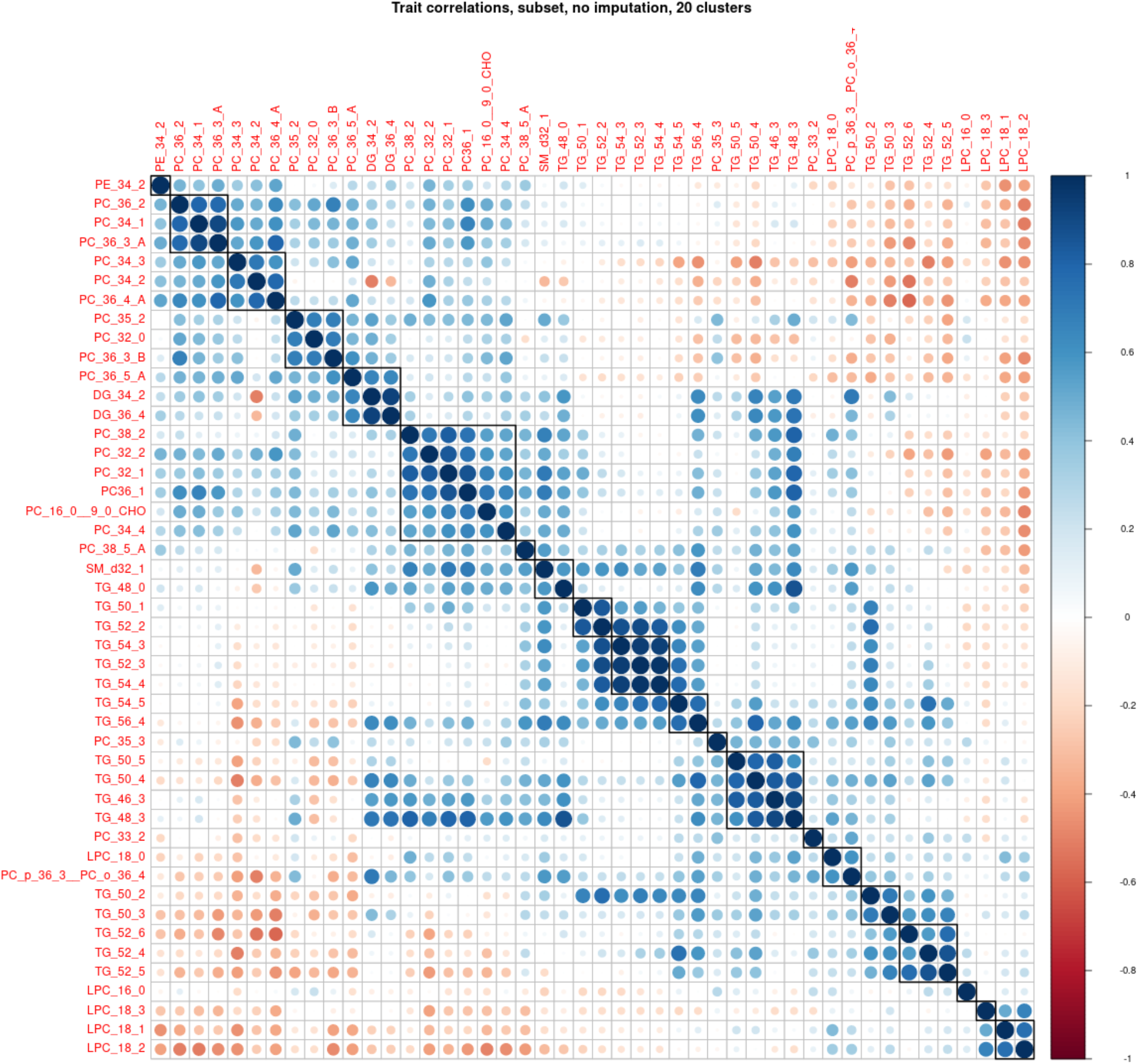
Cluster analysis of lipidomics data. Representative species from each cluster ere used for *Q_ST_* - *F_ST_* analysis

**Fig. S2.**
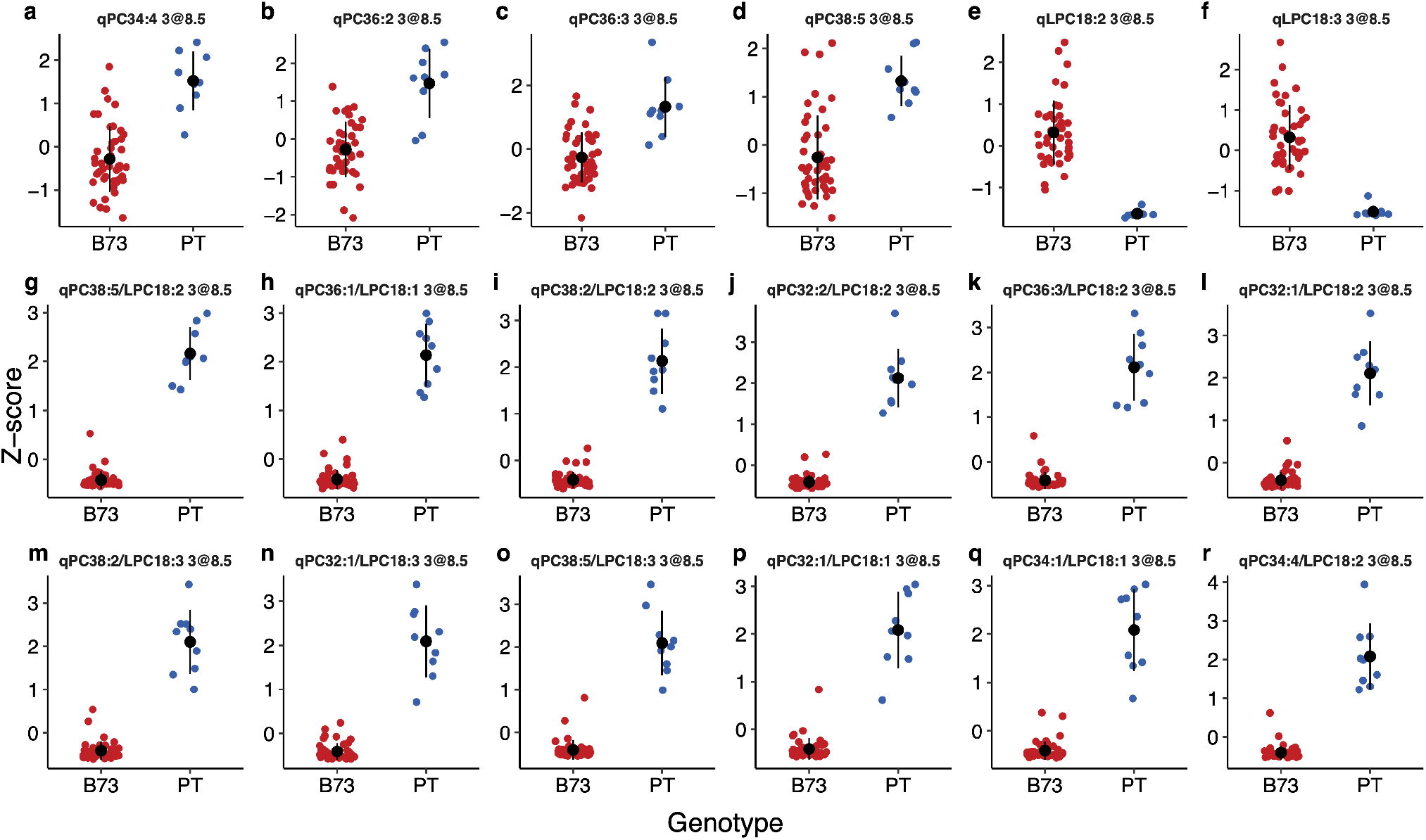
Effects of QTLs at chromosome 3 @8.5 Mb. **(A-D)** Effect for individual PC **(E-F)** and LPC species **(G-R)** The 12 most significant peaks for PC/LPC ratios

**Fig. S3.**
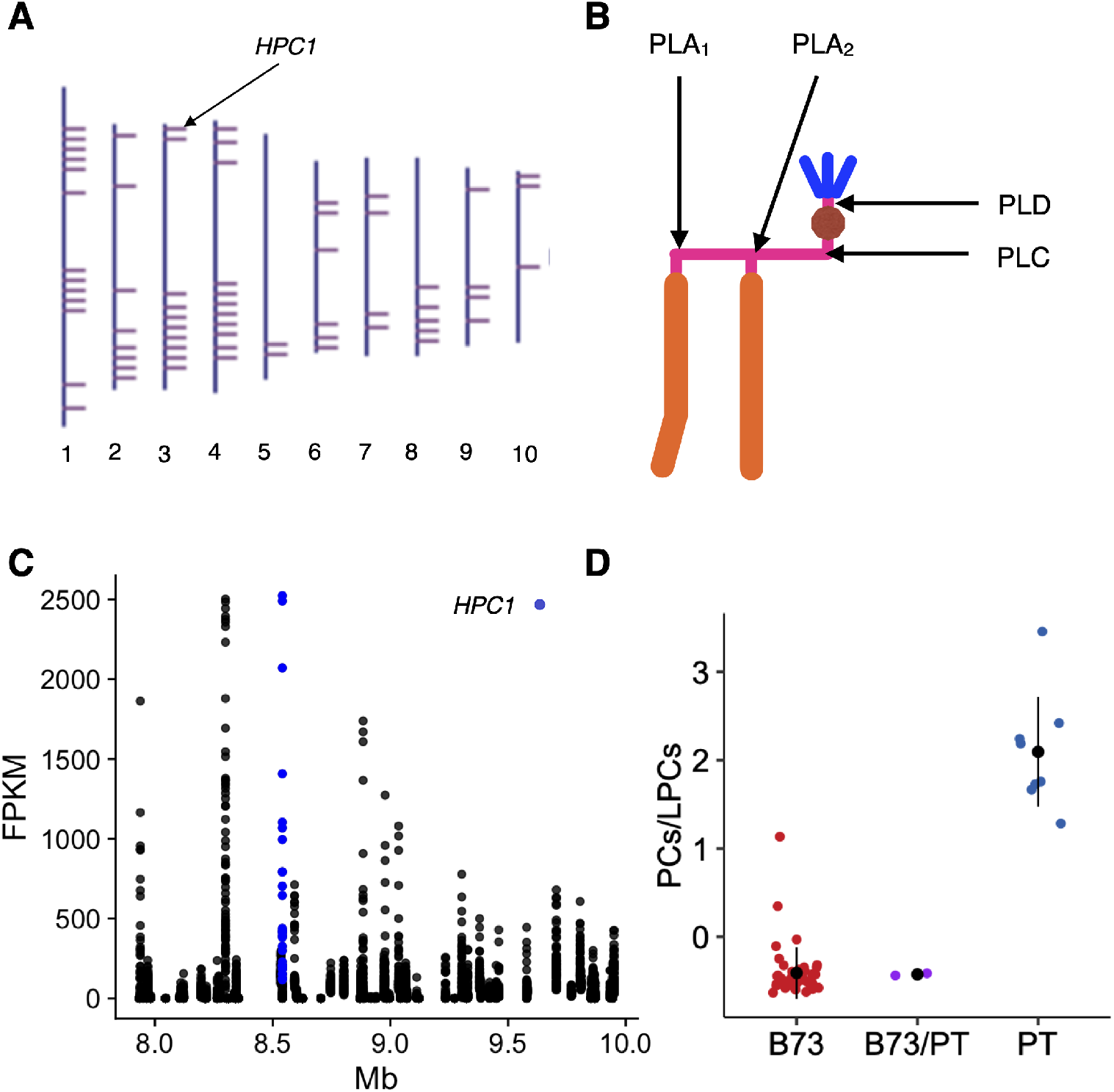
Phospholipase activity in maize. **A)** Genomic location of genes encoding proteins with predicted phospholipase A1 activity. **B)** Site of action of the different types of phospholipases. **C)** Expression levels from B73 leaves for genes in the 7.9–10 Mb QTL interval of chromosome 3. Data from (52) **D)** Effect sizes of PC/LPC levels at RILs homozyzogous for B73, PT or heterozygous at the QTL qPC/LPC3@8.5.

**Fig. S4.**
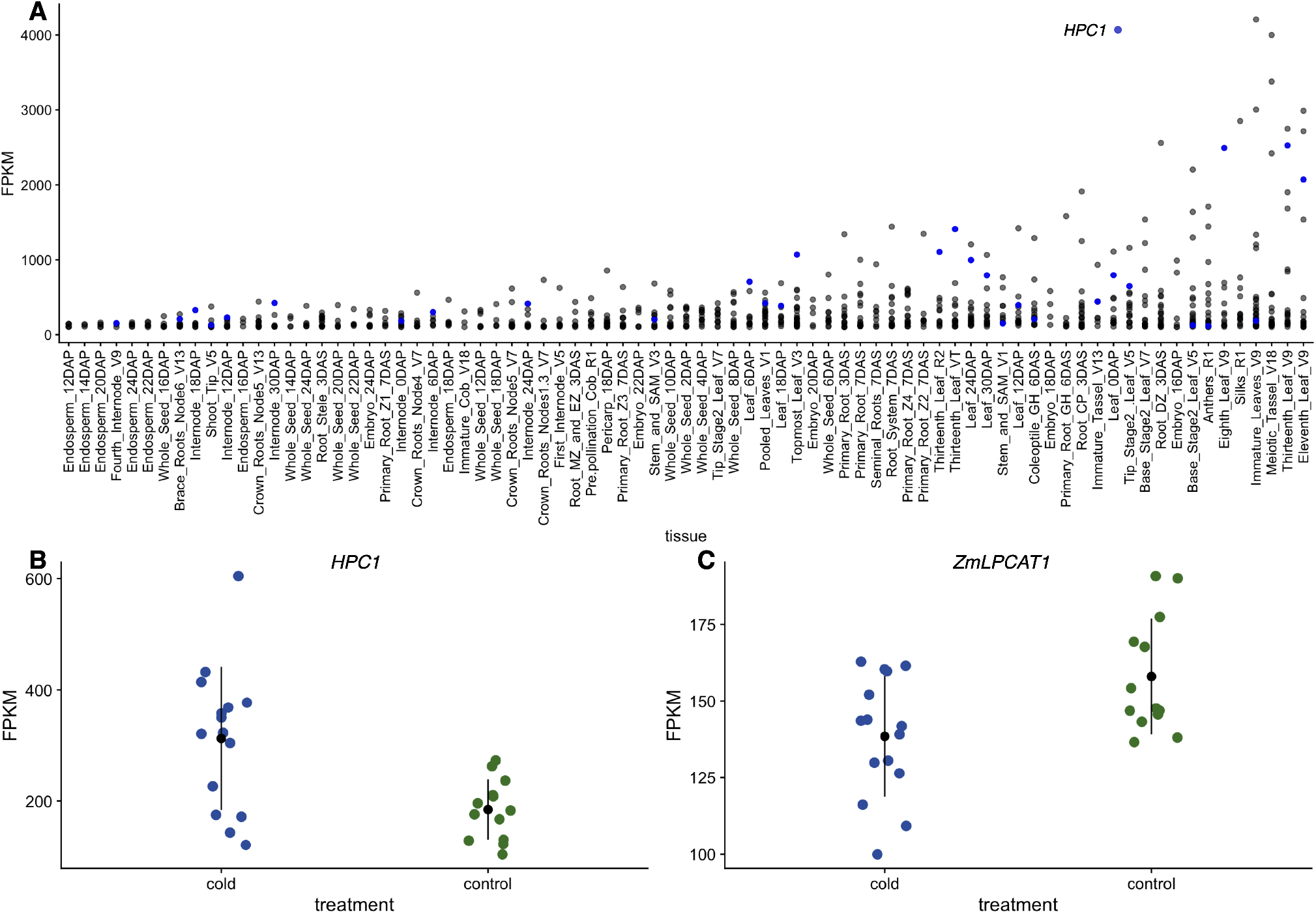
Phospholipase expression in temperate inbred lines. **A)** Expression levels in B73 of genes encoding enzymes with predicted phospholipase A1 activity across different tissues. is indicated in blue. Data from (52). **B)** and **C)**and *ZmLPCAT1* expression levels in the temperate inbred lines B73, Mo17, OH43, and PH207 under cold and control conditions. Values taken from (53)).

**Fig. S5.**
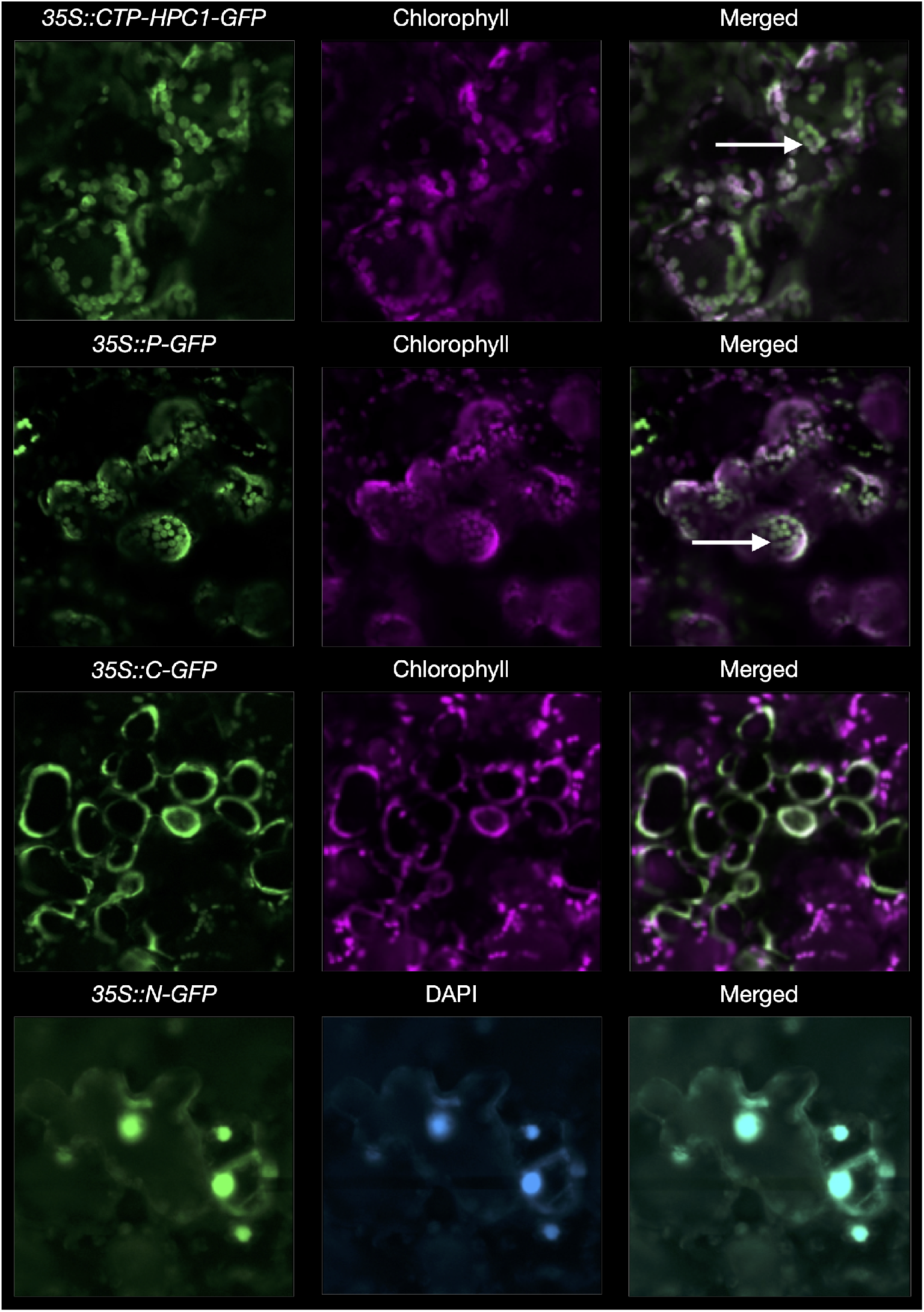
Subcellular localization of HPC1 in *Nicotiana benthamiana* epidermal leaf cells. CTP.HPC1-GFP, corresponding to the first 52 amino acids HPC1, were fused to GFP. Three constructs for various subcellular compartments were used as control; cytoplasm (C-GFP), nucleus (N-GFP), and chloroplast (P-GFP). All reporters were under the control of the cauliflower Mosaic Virus (CaMV) 35S promoter. The white arrows show chloroplast and nucleus.

**Fig. S6.**
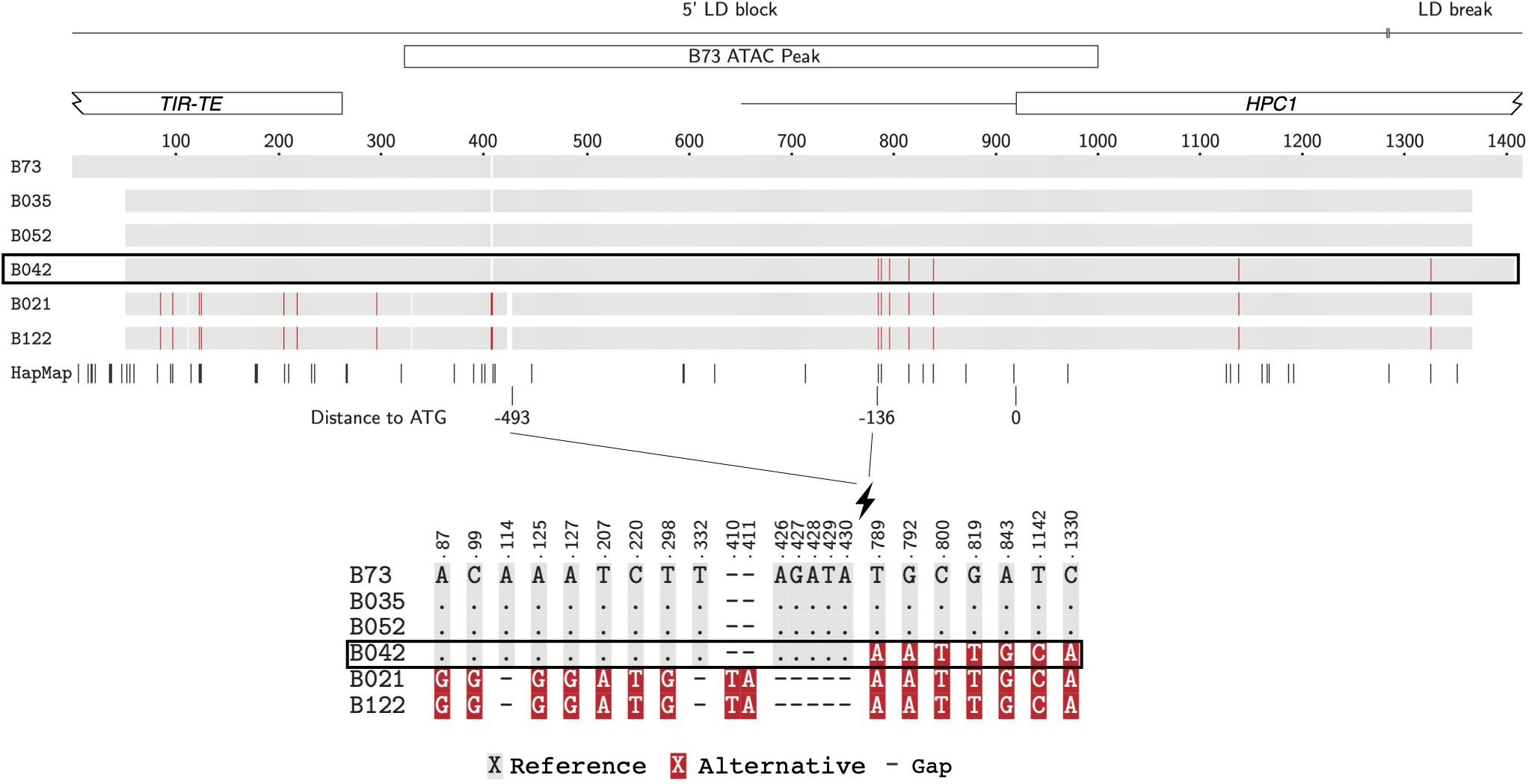
Recombination at the promoter in the B73 x PT segregating population. *Overview (top)* 1419 bp global alignment of Sanger sequences from B73 and 5 RILs (B035, B052, B042, B021 and B122). At variant positions B042 shows B73 genotype (grey) up to alignment position 430, and PT genotype (red) from 789 downstream. We infer that there is a recombination breakpoint between −493 and −136 bp upstream the translation start codon of B042. HapMap track shows HapMap2.7 SNPs used for Linkage Disequilibrium calculations in MaizeSNPDB as in Figure 5. The 5’ LD block spans the upstream intergenic region, including two ATAC-seq chromatin accessibility peaks (one included in this fragment), and up to 364 bp of coding sequence. Then there is a LD break, and a downstream 3’ LD block (not included in these sequences). *TIR-TE* terminal inverted repeat transposable element. B73 ATAC-seq: narrow peak showing chromatin accessibility in leaf tissue (54). *Base pair detail (bottom)* B73 and and PT nucleotide alleles are marked in grey and red respectively. Coordinate numbers on top of both views are global alignment positions.

**Fig. S7.**
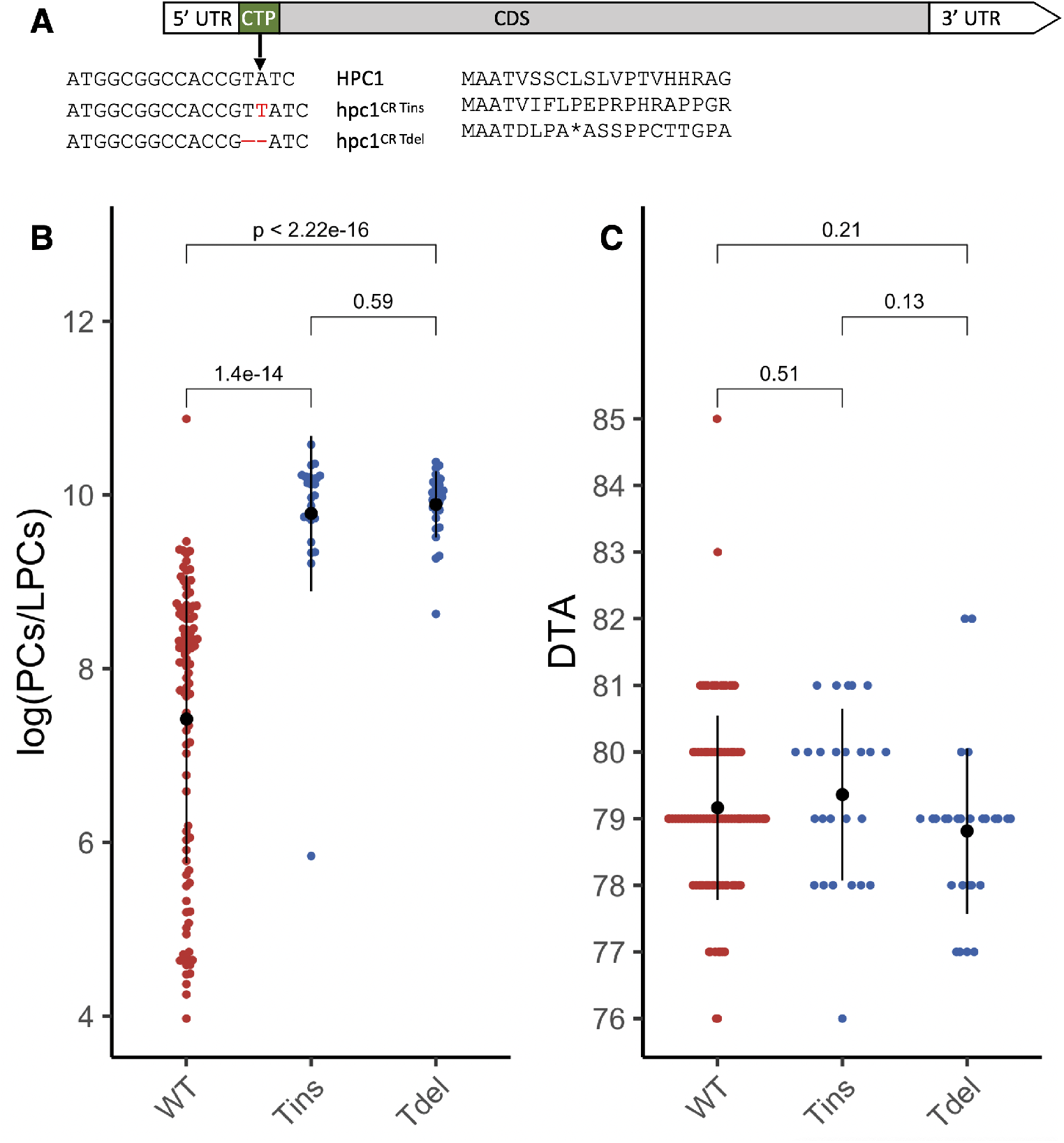
Effects of the *hpc1^CR^* allele. **A** Physical location of CRISPR mutant alleles. **B** log(PCs/LPCs) ratios of WT and the CRISPR mutant alleles. **C** Days to anthesis of the same plants shown in panel A.

**Fig. S8.**
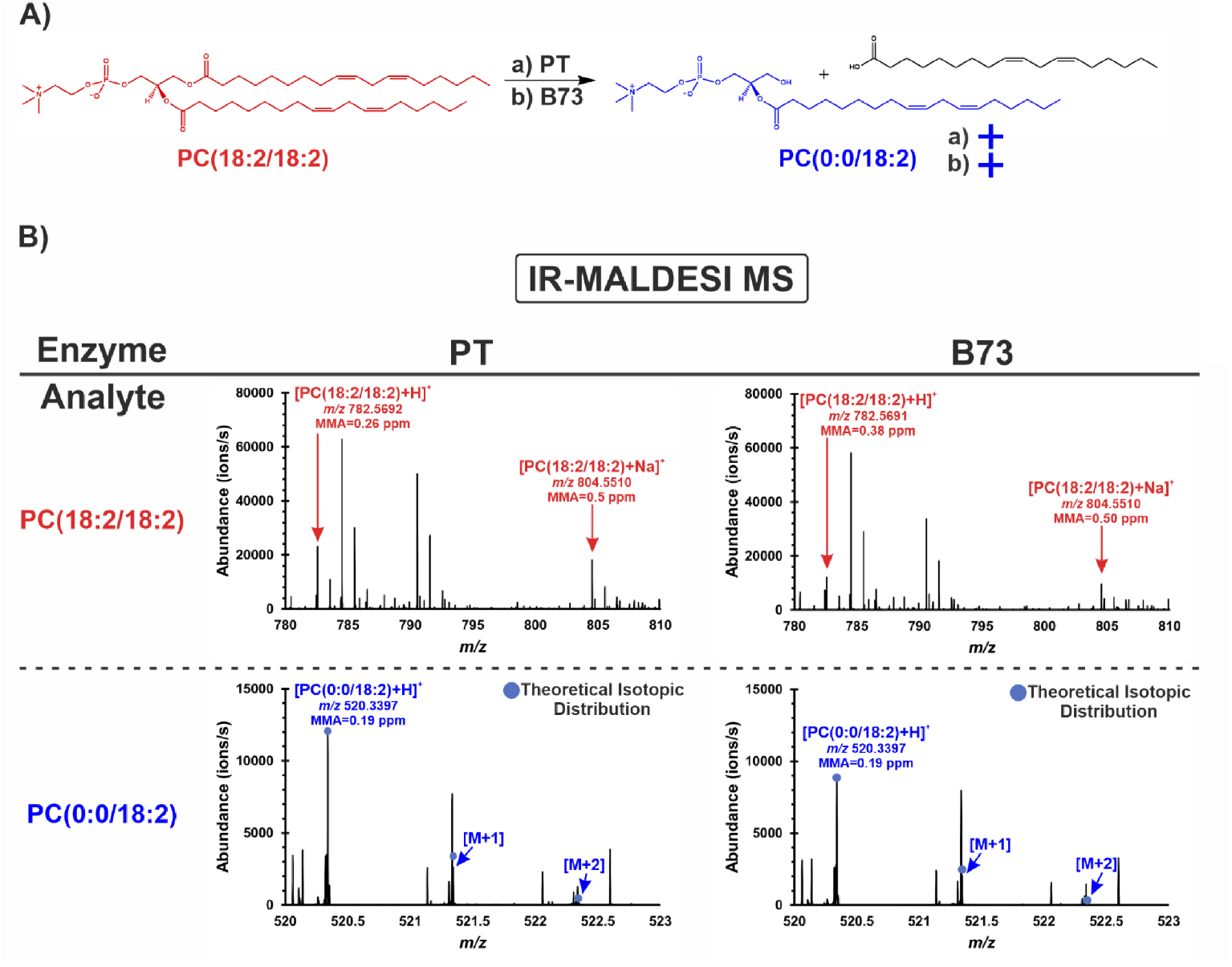
alleles have phospholipase activity. **A** Schematic of the reaction of PC(18:2/18:2) (red) to PC(0:0/18:2) (blue). **B** IR-MALDESI-MS mass spectra of the HPC1 reaction using the *HPC1-PT* and *HPC1-B73* alleles. The experimental *m/z* and mass measurement accuracy (MMA) is labelled below each identification. Both protonated and sodiated adducts of PC(18:2/18:2) were measured. Each mass spectrum was averaged over 100 scans.

**Fig. S9.**
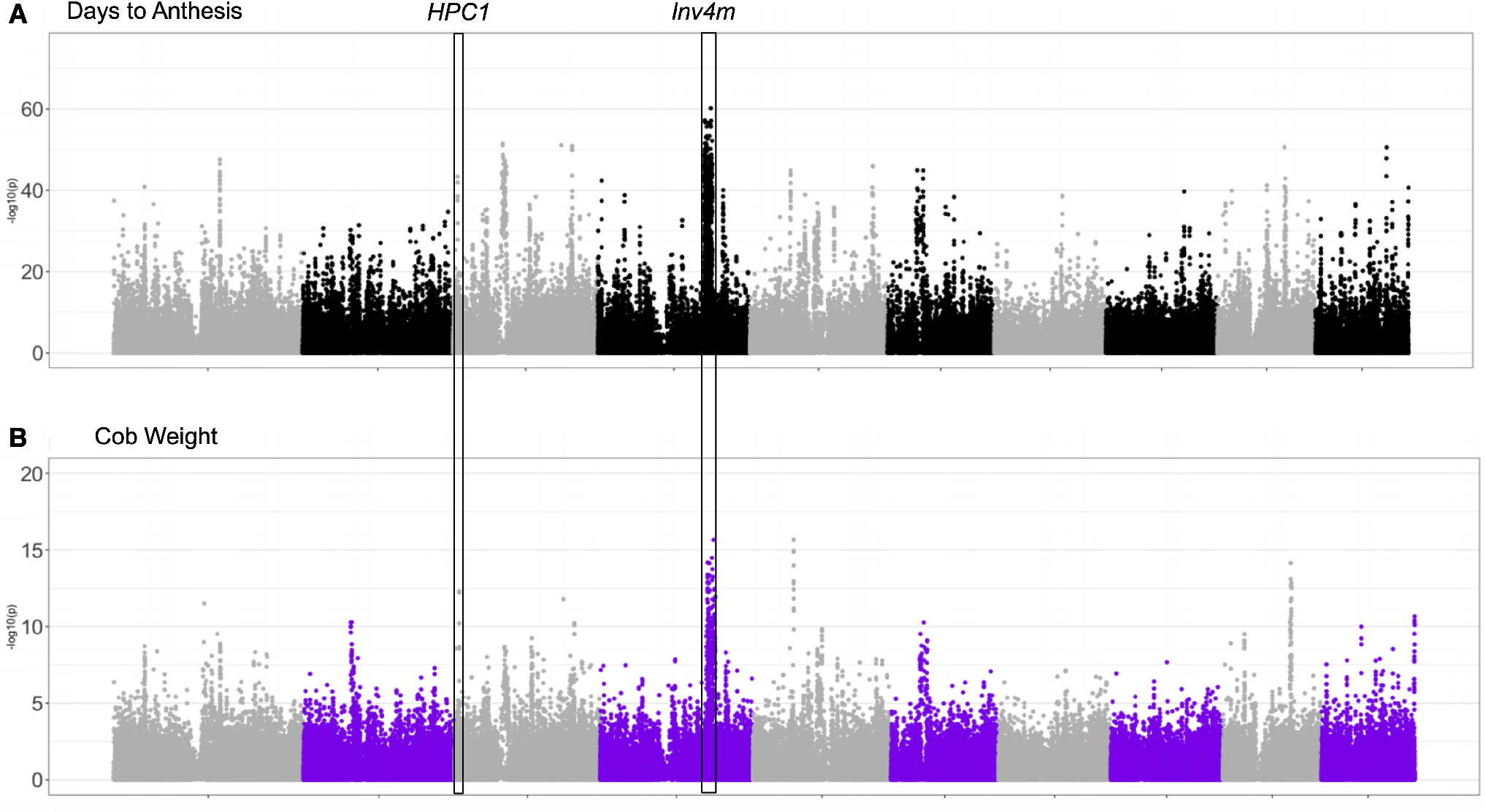
Genome wide association of genotype by elevation by genotype interactions of days to anthesis (**A**) and cob weight (**B**) using data from (3)

**Fig. S10.**
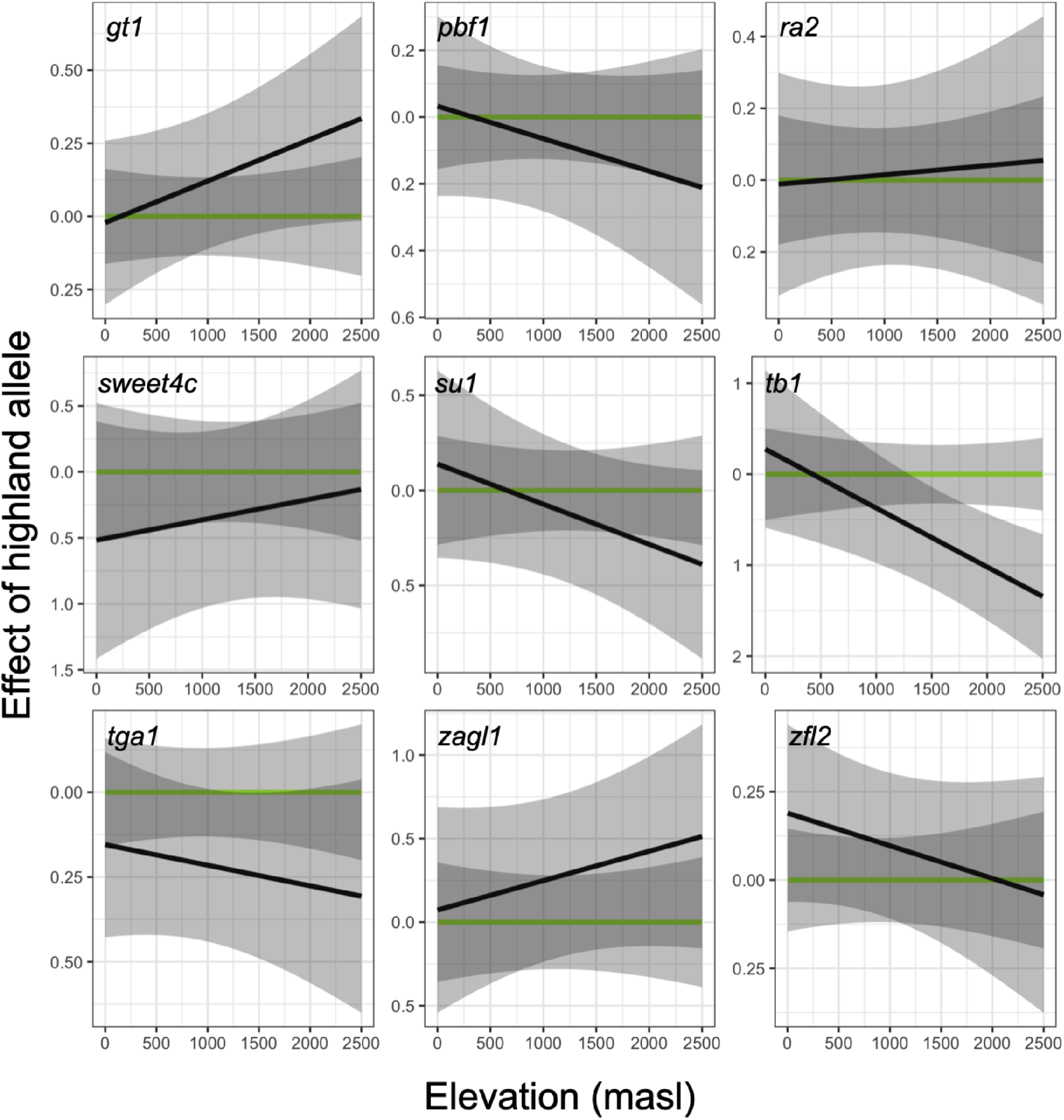
Effect of highland allele of several domestication and adaptation genes across different elevations.

**Fig. S11.**
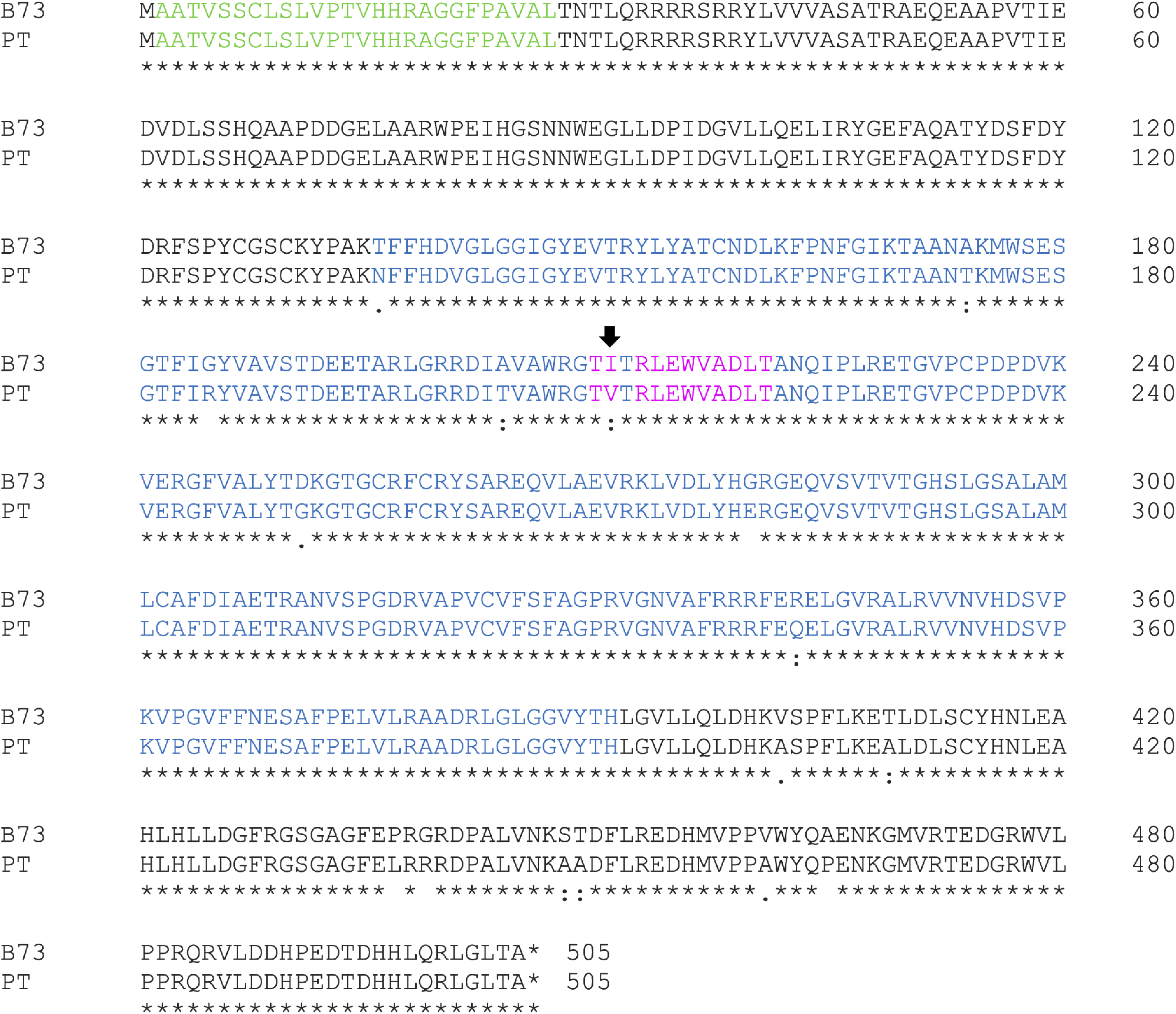
Alignment of HPC1-B73 and HPC1-PT using ClustalOmega. Green residues are those from the chloroplast transit peptide. Blue resides are part of the Lipase3 domain. Magenta residues are the flap lid. Arrow denotes I211V mutation. “*” denotes matching sequence, “:” denotes conservation between groups with similar properties, “.” denotes conservation of groups with weakly similar properties. . figure:Sup:aa*_a_lilnment*

**Fig. S12.**
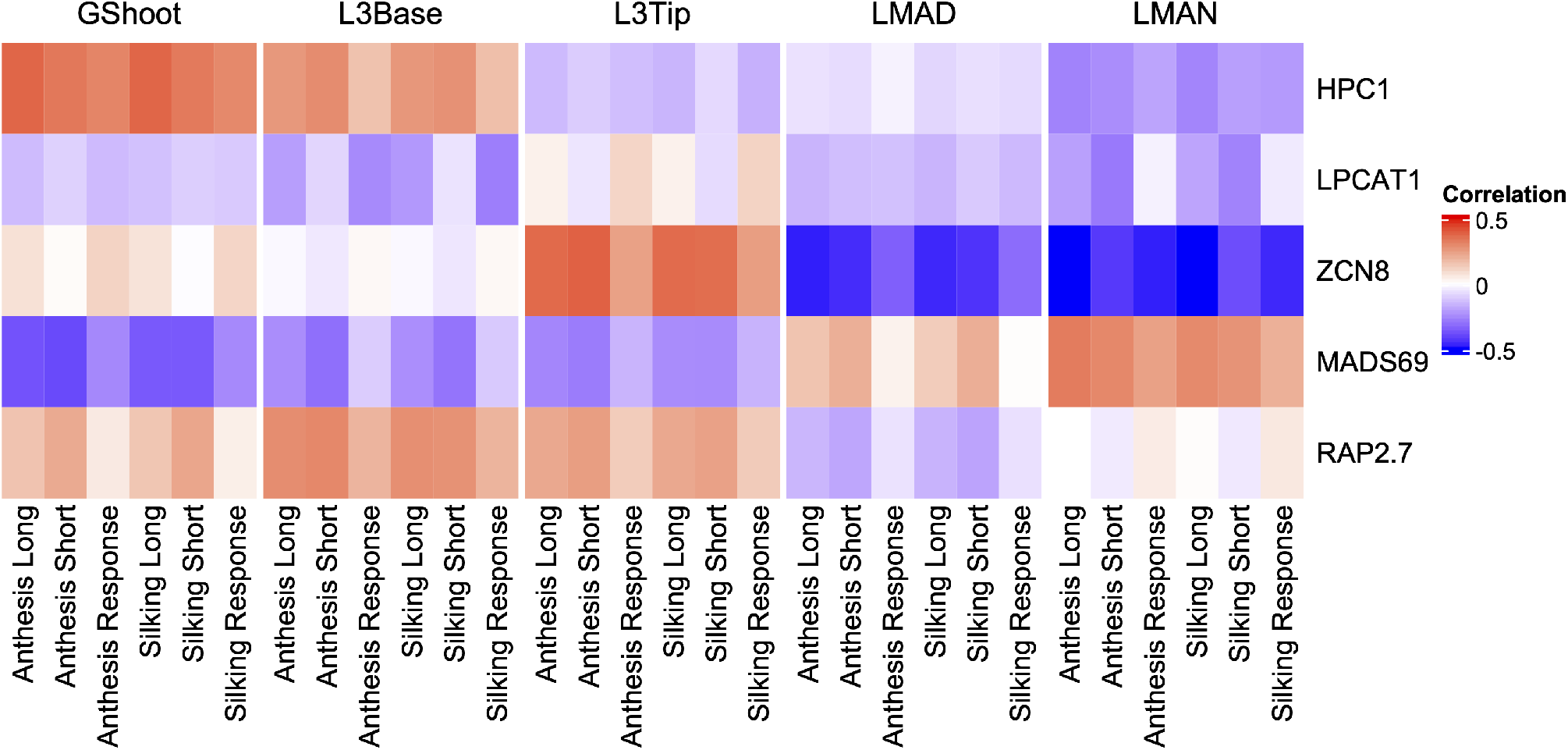
Correlation between flowering time and expression of phospholipid-related genes. Gene expression was correlated to flowering time traits from aerial tissues in the 282 diversity panel. Data obtained from (50).

**Fig. S13.**
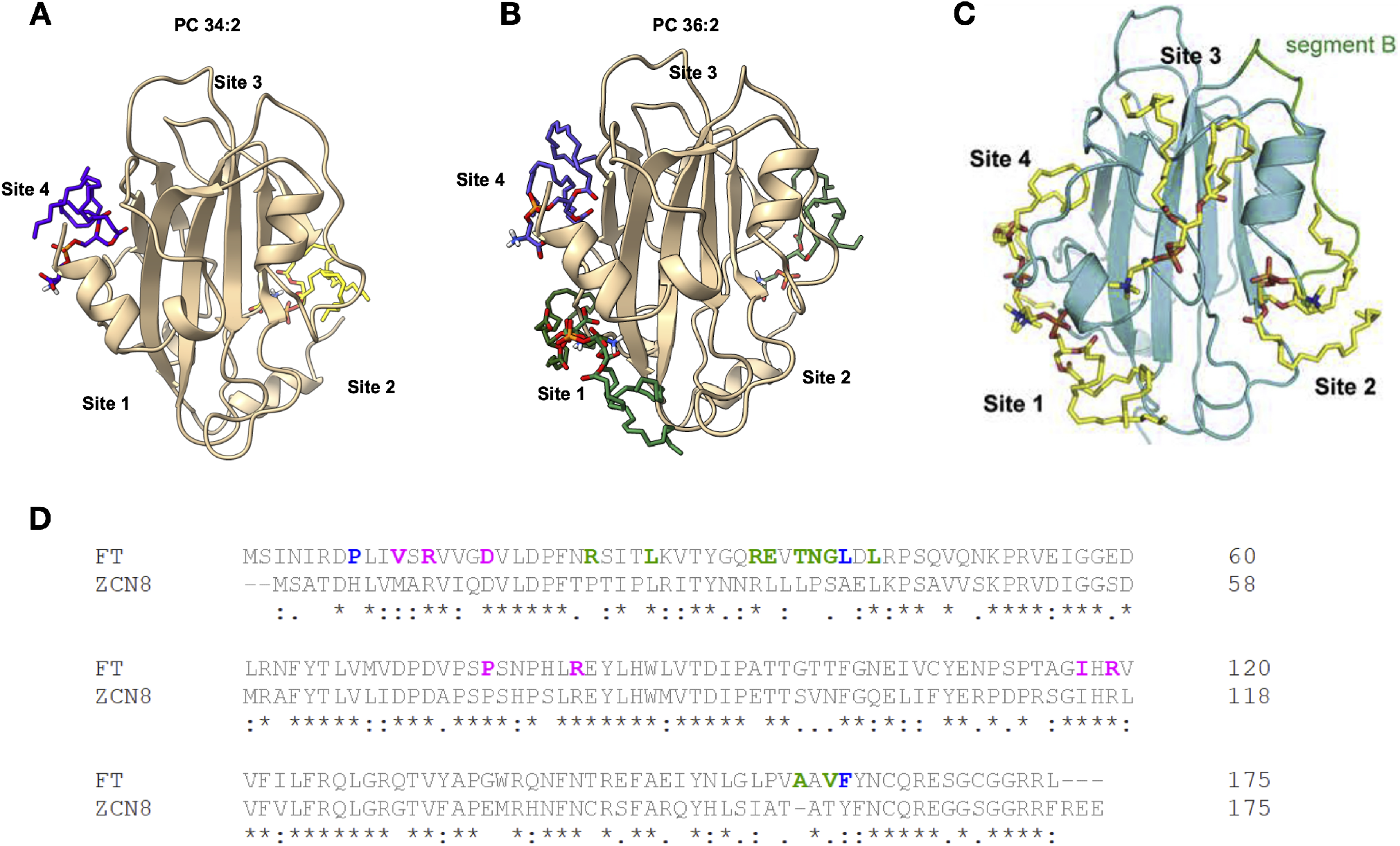
Predicted binding sites of ZCN8 align with binding sites of Arabidopsis FT. Shown are AutoDock Vina (55) ZCN8 - lipid docking interactions of a RoseTTAFold (56) model of ZCN8 PC 34:2 (**A**) and PC 36:2 (**B**) compared with the docking model of PC 36:2 *Arabidopsis thaliana* FT (**C**) from (57). Docking was performed on an NMRBox server (58). (**D**) Alignment of *Arabidopsis thaliana* FT and *Zea mays* ZCN8. Residues in blue are present in both docking site 1 and docking site 4 identified in (57). Residues in magenta are present only in docking site 1 while residues in green are present only in docking site 4. Symbols below the residues indicate level of alignment where * denotes complete alignment, : represents a conserved substitution, and. represents a semi-conserved substitution.

**Fig. S14.**
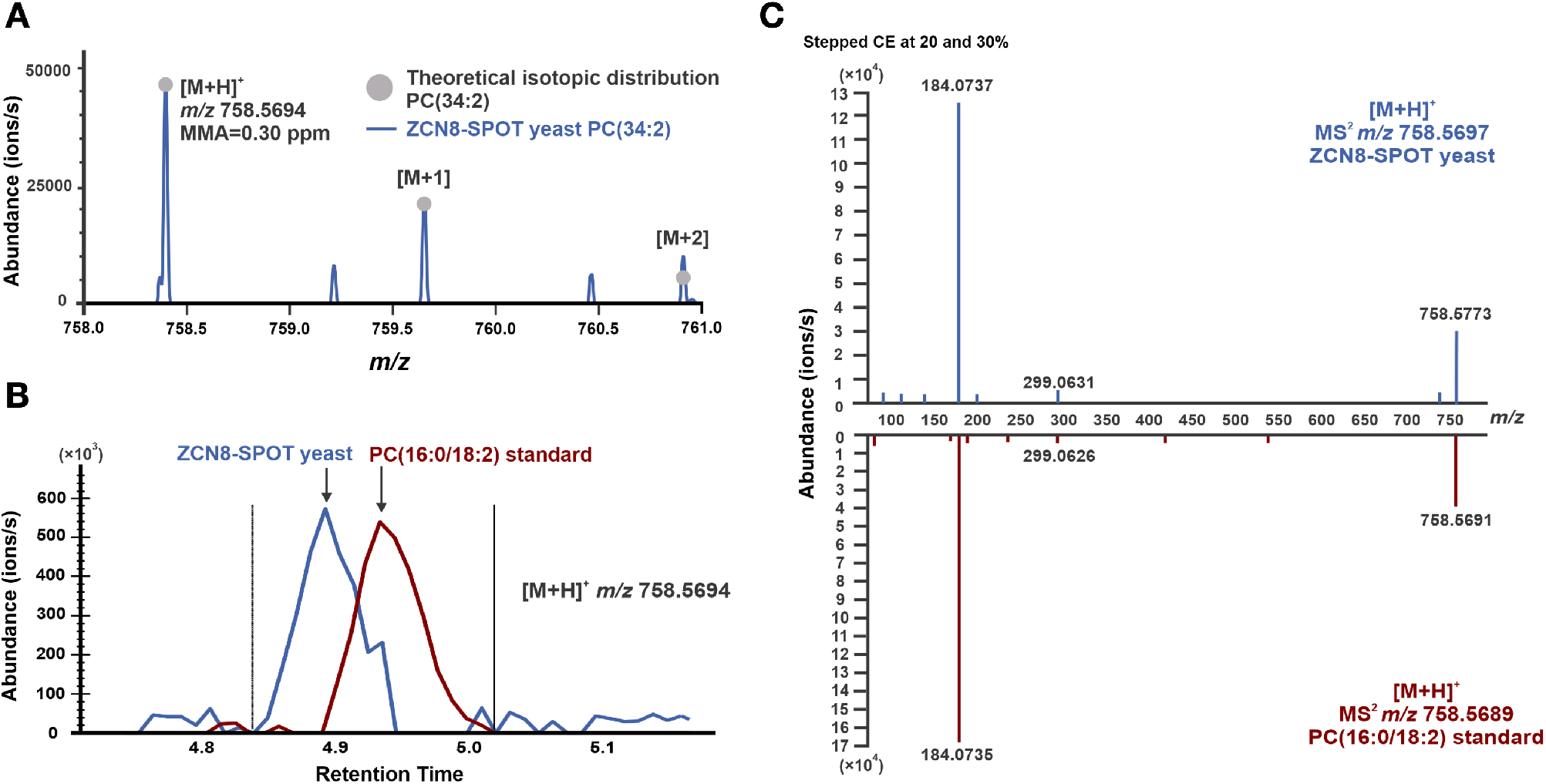
ZCN8 binds to phosphatidylcholine. **A** Mass spectrum of ZCN8-SPOT yeast sample compared to the theoretical isotopic pattern of PC(34:2) at [M+H]^+^ *m/z* 758.5694. The experimental *m/z* and mass measurement accuracy (MMA) are labeled. The spectrum of ZCN8-SPOT is an average of 16 scans across the chromatographic peak of *m/z* 758.5694. **B** Extracted ion chromatogram of *m/z* 758.5694 from the PC(16:0/18:2) standard and ZCN8-SPOT yeast from two separate injections. **C** MS^2^ fragmentation spectra comparison for *m/z* 758.5694 from ZCN8-SPOT yeast and the PC(16:0/18:2) standard. The comparison data between an authentic standard and ZCN8-SPOT yeast was acquired with the same lipid profiling method as described in the Materials and Methods section for the Thermo Scientific Orbitrap Exploris 480 mass spectrometer with the following modifications: the injection volume was 10 *μ*L, full scan spectra were acquired from *m/z* 200 – 1000 and *m/z* 758.5694 was included in the target mass list for MS^2^ selection.

## Footnotes

1 Open pollinated varieties that have been selected for specific uses and environments by smallholder farmers; they are characterized by similar morphotypes. Landraces typically show as much diversity within individuals from the same landrace group as between groups.

## References

[1] X Yi, et al., Sequencing of 50 human exomes reveals adaptation to high altitude. Science 329, 75–78 (2010).

[2] A Bigham, et al., Identifying signatures of natural selection in tibetan and andean populations using dense genome scan data. PLoS genetics 6, e1001116 (2010).

[3] J Yang, et al., Genetic signatures of high-altitude adaptation in tibetans. Proceedings of the National Academy of Sciences of the United States of America 114, 4189–4194 (2017).

[4] F Cicconardi, et al., Genomic signature of shifts in selection in a subalpine ant and its physiological adaptations (2020).

[5] EC Gilmore, Jr., JS Rogers, Heat units as a method of measuring maturity in corn 1. Agronomy journal 50, 611–615 (1958).

[6] JL Hatfield, JH Prueger, Temperature extremes: Effect on plant growth and development. Weather and Climate Extremes 10, 4–10 (2015).

[7] Y Matsuoka, et al., A single domestication for maize shown by multilocus microsatellite genotyping. Proceedings of the National Academy of Sciences of the United States of America 99, 6080–6084 (2002).

[8] DR Piperno, KV Flannery, The earliest archaeological maize (zea mays l.) from highland mexico: new accelerator mass spectrometry dates and their implications. Proceedings of the National Academy of Sciences of the United States of America 98, 2101–2103 (2001).

[9] MB Hufford, et al., The genomic signature of crop-wild introgression in maize. PLoS genetics 9, e1003477 (2013).

[10] E Gonzalez-Segovia, et al., Characterization of introgression from the teosinte zea mays ssp. mexicana to mexican highland maize. PeerJ 7, e6815 (2019).

[11] JA Aguirre-Liguori, et al., Divergence with gene flow is driven by local adaptation to temperature and soil phosphorus concentration in teosinte subspecies (zea mays parviglumis and zea mays mexicana). Molecular ecology 24, 2663 (2019).

[12] N Lauter, C Gustus, A Westerbergh, J Doebley, The inheritance and evolution of leaf pigmentation and pubescence in teosinte. Genetics 167, 1949–1959 (2004).

[13] AK Hardacre, HA Eagles, Comparisons among populations of maize for growth at 13°c. Crop science 20, 780–784 (1980).

[14] T Crow, et al., Gene regulatory effects of a large chromosomal inversion in highland maize. PLoS genetics 16, e1009213 (2020).

[15] DJ Gates, D Runcie, GM Janzen, AR Navarro, others, Singlegene resolution of locally adaptive genetic variation in mexican maize. bioRxiv (2019).

[16] J Stephen Athens, et al., Early prehistoric maize in northern highland ecuador. Latin American Antiquity 27, 3–21 (2016).

[17] A Grobman, et al., Preceramic maize from paredones and huaca prieta, peru. Proceedings of the National Academy of Sciences of the United States of America 109, 1755–1759 (2012).

[18] S Takuno, et al., Independent Molecular Basis of Convergent Highland Adaptation in Maize. Genetics, genet-ics.115.178327 (2015).

[19] L Wang, et al., Molecular parallelism underlies convergent highland adaptation of maize landraces. bioRxiv (2020).

[20] RR da Fonseca, et al., The origin and evolution of maize in the southwestern united states. Nature plants 1, 14003 (2015).

[21] K Swarts, et al., Genomic estimation of complex traits reveals ancient maize adaptation to temperate north america. Science 357, 512–515 (2017).

[22] HY Hung, et al., ZmCCT and the genetic basis of day-length adaptation underlying the postdomestication spread of maize (2012).

[23] Z Liang, et al., Conventional and hyperspectral time-series imaging of maize lines widely used in field trials. Giga-Science 7, 1–11 (2018).

[24] L Guo, et al., Stepwise cis-regulatory changes in ZCN8 contribute to maize Flowering-Time adaptation. Current biology: CB 28, 3005–3015.e4 (2018).

[25] ND Coles, MD McMullen, PJ Balint-Kurti, RC Pratt, JB Holland, Genetic control of photoperiod sensitivity in maize revealed by joint multiple population analysis. Genetics 184, 799–812 (2010).

[26] C Huang, et al., ZmCCT9 enhances maize adaptation to higher latitudes. Proceedings of the National Academy of Sciences of the United States of America 115, E334–E341 (2018).

[27] Q Yang, et al., CACTA-like transposable element in Zm-CCT attenuated photoperiod sensitivity and accelerated the postdomestication spread of maize. Proceedings of the National Academy of Sciences of the United States of America 110, 16969–16974 (2013).

[28] S Salvi, et al., Conserved noncoding genomic sequences associated with a flowering-time quantitative trait locus in maize. Proceedings of the National Academy of Sciences of the United States of America 104, 11376–11381 (2007).

[29] JT Brandenburg, et al., Independent introductions and admixtures have contributed to adaptation of european maize and its american counterparts. PLoS genetics 13, e1006666 (2017).

[30] T Degenkolbe, et al., Differential remodeling of the lipidome during cold acclimation in natural accessions of arabidopsis thaliana. The Plant journal: for cell and molecular biology 72, 972–982 (2012).

[31] R Welti, et al., Profiling membrane lipids in plant stress responses: Role of phospholipase D*α* in freezing-induced lipid changes in arabidopsis. The Journal of biological chemistry 277, 31994–32002 (2002).

[32] DV Lynch, PL Steponkus, Plasma membrane lipid alterations associated with cold acclimation of winter rye seedlings (secale cereale l. cv puma). Plant physiology 83, 761–767 (1987).

[33] J Jouhet, Importance of the hexagonal lipid phase in biological membrane organization. Front. Plant Sci. 4, 494 (2013).

[34] JR Henriksen, TL Andresen, LN Feldborg, L Duelund, JH Ipsen, Understanding detergent effects on lipid membranes: a model study of lysolipids. Biophys. J. 98, 2199–2205 (2010).

[35] L Yan, et al., Parallels between natural selection in the cold-adapted crop-wild relative tripsacum dactyloides and artificial selection in temperate adapted maize. The Plant journal: for cell and molecular biology (2019).

[36] H Susila, et al., Florigen sequestration in cellular membranes modulates temperature-responsive flowering. Science 373, 1137–1142 (2021).

[37] Y Gu, et al., Biochemical and transcriptional regulation of membrane lipid metabolism in maize leaves under low temperature. Front. Plant Sci. 8, 2053 (2017).

[38] Y Nakamura, et al., Arabidopsis florigen FT binds to diurnally oscillating phospholipids that accelerate flowering. Nature communications 5, 3553 (2014).

[39] C Riedelsheimer, Y Brotman, M Méret, AE Melchinger, L Willmitzer, The maize leaf lipidome shows multilevel genetic control and high predictive value for agronomic traits. Scientific reports 3, 2479 (2013).

[40] T Leinonen, RJS McCairns, RB O’Hara, J Merilä, Q(ST)-F(ST) comparisons: evolutionary and ecological insights from genomic heterogeneity. Nature reviews. Genetics 14, 179–190 (2013).

[41] GM Janzen, et al., Demonstration of local adaptation of maize landraces by reciprocal transplantation. (2021).

[42] JE Pool, DT Braun, JB Lack, Parallel evolution of cold tolerance within drosophila melanogaster. Molecular biology and evolution 34, 349–360 (2017).

[43] S Yeaman, AC Gerstein, KA Hodgins, MC Whitlock, Quantifying how constraints limit the diversity of viable routes to adaptation. PLOS Genetics 14, e1007717 (2018) Publisher: Public Library of Science.

[44] JA Romero Navarro, et al., A study of allelic diversity underlying flowering-time adaptation in maize landraces. Nature genetics 49, 476 (2017).

[45] K Luu, E Bazin, MGB Blum, pcadapt: an R package to perform genome scans for selection based on principal component analysis. Molecular ecology resources 17, 67–77 (2017).

[46] L Wang, et al., Metabolic Interactions between the Lands Cycle and the Kennedy Pathway of Glycerolipid Synthesis in *Arabidopsis* Developing Seeds. The Plant Cell 24, 4652–4669 (2012).

[47] GS Richmond, TK Smith, Phospholipases A1. International Journal of Molecular Sciences 12, 588–612 (2011).

[48] K Wang, et al., Two Abscisic Acid-Responsive Plastid Lipase Genes Involved in Jasmonic Acid Biosynthesis in *Arabidopsis thaliana*. The Plant Cell 30, 1006–1022 (2018).

[49] S Ishiguro, A Kawai-Oda, J Ueda, I Nishida, K Okada, The DEFECTIVE IN ANTHER DEHISCIENCE gene encodes a novel phospholipase A1 catalyzing the initial step of jasmonic acid biosynthesis, which synchronizes pollen maturation, anther dehiscence, and flower opening in arabidopsis. Plant Cell 13, 2191–2209 (2001).

[50] S Perez-Limón, et al., A B73 × Palomero toluqueño mapping population reveals local adaptation in mexican highland maize. G3 GeneslGenomeslGenetics (2022).

[51] SE Jensen, LC Johnson, T Casstevens, ES Buckler, Predicting protein domain temperature adaptation across the prokaryote-eukaryote divide. (2021).

[52] SE Jensen, ES Buckler, Pfam domain adaptation profiles reflect plant species, evolutionary history. (2021).

[53] KW Broman, H Wu, S Sen, GA Churchill, R/qtl: QTL mapping in experimental crosses. Bioinformatics 19, 889–890 (2003).

[54] SB Ryu, Phospholipid-derived signaling mediated by phospholipase a in plants. Trends in plant science 9, 229–235 (2004).

[55] SC Stelpflug, et al., An Expanded Maize Gene Expression Atlas based on RNA Sequencing and its Use to Explore Root Development. The plant genome 9, 0 (2016).

[56] O Emanuelsson, H Nielsen, G von Heijne, ChloroP, a neural network-based method for predicting chloroplast transit peptides and their cleavage sites. Protein Sci. 8, 978–984 (1999).

[57] DE Runcie, L Crawford, Fast and flexible linear mixed models for genome-wide genetics. PLOS Genetics 15, 1–24 (2019).

[58] W Zhou, L Wang, W Zheng, W Yao, MaizeSNPDB: A comprehensive database for efficient retrieve and analysis of SNPs among 1210 maize lines. Computational and Structural Biotechnology Journal 17, 1377–1383 (2019).

[59] WA Ricci, et al., Widespread long-range cis-regulatory elements in the maize genome. Nature plants 5, 1237–1249 (2019).

[60] JG Wallace, et al., Association mapping across numerous traits reveals patterns of functional variation in maize. PLoS genetics 10, e1004845 (2014).

[61] MR Woodhouse, et al., A pan-genomic approach to genome databases using maize as a model system. BMC plant biology 21, 385 (2021).

[62] SC Potter, et al., HMMER web server: 2018 update (2018).

[63] S El-Gebali, et al., The pfam protein families database in 2019. Nucleic acids research 47, D427–D432 (2019).

[64] E Cimen, SE Jensen, ES Buckler, Building a tRNA thermometer to estimate microbial adaptation to temperature. Nucleic Acids Research (2020).

[65] KAG Kremling, et al., Dysregulation of expression correlates with rare-allele burden and fitness loss in maize. Nature 555, 520–523 (2018).

[66] R Bukowski, et al., Construction of the third generation zea mays haplotype map. GigaScience (2017).

[67] Y Nakamura, et al., High-Resolution crystal structure of arabidopsis FLOWERING LOCUS T illuminates its Phospholipid-Binding site in flowering. iScience 21, 577–586 (2019).

[68] Y Nakamura, Plant phospholipid diversity: Emerging functions in metabolism and Protein–Lipid interactions. Trends in plant science 22, 1027–1040 (2017).

[69] EJ Veneklaas, et al., Opportunities for improving phosphorus-use efficiency in crop plants: Tansley review. The New phytologist 195, 306–320 (2012).

[70] A Cruz-Ramirez, et al., The xipotl mutant of Arabidopsis reveals a critical role for phospholipid metabolism in root system development and epidermal cell integrity. The Plant cell 16, 2020–2034 (2004).

[71] H Lambers, et al., Proteaceae from severely phosphorus-impoverished soils extensively replace phospholipids with galactolipids and sulfolipids during leaf development to achieve a high photosynthetic phosphorus-use-efficiency. The New phytologist 196, 1098–1108 (2012).

[72] SR Marla, et al., Comparative transcriptome and lipidome analyses reveal molecular chilling responses in Chilling-Tolerant sorghums. Plant Genome 10 (2017).

[73] L Qu, YJ Chu, WH Lin, HW Xue, A secretory phospholipase D hydrolyzes phosphatidylcholine to suppress rice heading time. PLoS genetics 17, e1009905 (2021).

[74] KL Mercer, H Perales, Structure of local adaptation across the landscape: flowering time and fitness in mexican maize (zea mays l. subsp. mays) landraces. Genetic resources and crop evolution 66, 27–45 (2019).

[75] AJ Waters, et al., Natural variation for gene expression responses to abiotic stress in maize. The Plant Journal 89, 706–717 (2017).

[76] AK Hottes, et al., Bacterial adaptation through loss of function. PLoS genetics 9, e1003617 (2013).

[77] FI Khan, et al., The lid domain in lipases: Structural and functional determinant of enzymatic properties. Frontiers in bioengineering and biotechnology 5, 16 (2017).

[78] Y Kim, R Nielsen, Linkage disequilibrium as a signature of selective sweeps. Genetics 167, 1513–1524 (2004).

[79] S Khan, SC Rowe, FG Harmon, Coordination of the maize transcriptome by a conserved circadian clock. BMC Plant Biol. 10, 126 (2010).

[80] S Maatta, et al., Levels of arabidopsis thaliana leaf phosphatidic acids, phosphatidylserines, and most Trienoate-Containing polar lipid molecular species increase during the dark period of the diurnal cycle. Front. Plant Sci. 3, 49 (2012).

[81] E Calfee, et al., Selective sorting of ancestral introgression in maize and teosinte along an elevational cline. PLoS genetics 17, e1009810 (2021).

[82] CM Lazakis, V Coneva, J Colasanti, ZCN8 encodes a potential orthologue of arabidopsis FT florigen that integrates both endogenous and photoperiod flowering signals in maize. J. Exp. Bot. 62, 4833–4842 (2011).

[83] T Guo, et al., Optimal designs for genomic selection in hybrid crops. Molecular plant 12, 390–401 (2019).

[84] O Trott, AJ Olson, AutoDock vina: improving the speed and accuracy of docking with a new scoring function, efficient optimization, and multithreading. Journal of computational chemistry 31, 455–461 (2010).

[85] M Baek, et al., Accurate prediction of protein structures and interactions using a three-track neural network. Science 373, 871–876 (2021).

[86] MW Maciejewski, et al., Nmrbox: A resource for biomolecular nmr computation. Biophysical Journal 112, 1529–1534 (2017).

## References

1. L Wang, et al., Molecular parallelism underlies convergent highland adaptation of maize landraces. bioRxiv (2020).

2. L Wang, et al., The interplay of demography and selection during maize domestication and expansion. Genome biology 18, 215 (2017).

3. DJ Gates, D Runcie, GM Janzen, AR Navarro, others, Single-gene resolution of locally adaptive genetic variation in mexican maize. bioRxiv (2019).

4. K Luu, E Bazin, MGB Blum, pcadapt: an R package to perform genome scans for selection based on principal component analysis. Mol. ecology resources 17, 67–77 (2017).

5. V Matyash, G Liebisch, TV Kurzchalia, A Shevchenko, D Schwudke, Lipid extraction by methyl-tert-butyl ether for high-throughput lipidomics. J. lipid research 49, 1137–1146 (2008).

6. H Tsugawa, et al., MS-DIAL: data-independent MS/MS deconvolution for comprehensive metabolome analysis. Nat. methods 12, 523–526 (2015).

7. S Fan, et al., Systematic error removal using random forest for normalizing large-scale untargeted lipidomics data. Anal. Chem. 91, 3590–3596 (2019) PMID: 30758187.

8. BC DeFelice, et al., Mass spectral feature list optimizer (ms-flo): A tool to minimize false positive peak reports in untargeted liquid chromatography–mass spectroscopy (lc-ms) data processing. Anal. Chem. 89, 3250–3255 (2017) PMID: 28225594.

9. KJ Adams, et al., Skyline for small molecules: A unifying software package for quantitative metabolomics. J. Proteome Res. 19, 1447–1458 (2020).

10. MC Whitlock, Evolutionary inference from qst. Mol. ecology 17, 1885–1896 (2008).

11. GM Janzen, et al., Demonstration of local adaptation of maize landraces by reciprocal transplantation. (2021).

12. PC Bürkner, brms: An r package for bayesian multilevel models using stan. J. statistical software 80, 1–28 (2017).

13. G Covarrubias-Pazaran, Genome-assisted prediction of quantitative traits using the r package sommer. PloS one 11, e0156744 (2016).

14. T Leinonen, RJS McCairns, RB O’Hara, J Merilä, Q(ST)-F(ST) comparisons: evolutionary and ecological insights from genomic heterogeneity. Nat. reviews. Genet. 14, 179–190 (2013).

15. B Gruber, PJ Unmack, OF Berry, A Georges, dartr: an r package to facilitate analysis of snp data generated from reduced representation genome sequencing. Mol. ecology resources 18, 691–699 (2018).

16. N Duforet-Frebourg, E Bazin, MG Blum, Genome scans for detecting footprints of local adaptation using a bayesian factor model. Mol. biology evolution 31, 2483–2495 (2014).

17. M Kanehisa, Y Sato, M Furumichi, K Morishima, M Tanabe, New approach for understanding genome variations in KEGG. Nucleic Acids Res. 47, D590–D595 (2019) Publisher: Oxford Academic.

18. JL Portwood, et al., MaizeGDB 2018: the maize multi-genome genetics and genomics database. Nucleic Acids Res. 47, D1146–D1154 (2019) Publisher: Oxford Academic.

19. JR Walsh, et al., The quality of metabolic pathway resources depends on initial enzymatic function assignments: a case for maize. BMC Syst. Biol. 10, 129 (2016).

20. FR Zapata, sawers-rellan-labs/pglipid: Glycerophospholipid Pathway analysis (2020).

21. S Yeaman, AC Gerstein, KA Hodgins, MC Whitlock, Quantifying how constraints limit the diversity of viable routes to adaptation. PLOS Genet. 14, e1007717 (2018) Publisher: Public Library of Science.

22. Q Wu, et al., The maize heterotrimeric G protein *—* subunit controls shoot meristem development and immune responses. Proc. Natl. Acad. Sci. U. S. A. 117, 1799–1805 (2020).

23. SN Char, et al., An agrobacterium-delivered CRISPR/Cas9 system for high-frequency targeted mutagenesis in maize. Plant Biotechnol. J. 15, 257–268 (2017).

24. VA Brazelton, Jr, et al., A quick guide to CRISPR sgRNA design tools. GM Crop. Food 6, 266–276 (2015).

25. M Caleb Bagley, KP Garrard, DC Muddiman, The development and application of matrix assisted laser desorption electrospray ionization: The teenage years. Mass Spectrom. Rev. (2021).

26. MT Bokhart, DC Muddiman, Infrared matrix-assisted laser desorption electrospray ionization mass spectrometry imaging analysis of biospecimens. Analyst 141, 5236–5245 (2016).

27. KP Garrard, M Ekelöf, S Khodjaniyazova, MC Bagley, DC Muddiman, A versatile platform for mass spectrometry imaging of arbitrary spatial patterns. J. Am. Soc. Mass Spectrom. 31, 2547–2552 (2020).

28. MC Chambers, et al., A cross-platform toolkit for mass spectrometry and proteomics. Nat. Biotechnol. 30, 918–920 (2012).

29. AM Race, IB Styles, J Bunch, Inclusive sharing of mass spectrometry imaging data requires a converter for all. J. Proteomics 75, 5111–5112 (2012).

30. G Robichaud, KP Garrard, JA Barry, DC Muddiman, MSiReader: an open-source interface to view and analyze high resolving power MS imaging files on matlab platform. J. Am. Soc. Mass Spectrom. 24, 718–721 (2013).

31. MT Bokhart, M Nazari, KP Garrard, DC Muddiman, MSiReader v1.0: Evolving Open-Source mass spectrometry imaging software for targeted and untargeted analyses. J. Am. Soc. Mass Spectrom. 29, 8–16 (2018).

32. T Crow, et al., Gene regulatory effects of a large chromosomal inversion in highland maize. PLoS genetics 16, e1009213 (2020).

33. BT Townsley, MF Covington, Y Ichihashi, K Zumstein, NR Sinha, Brad-seq: Breath adapter directional sequencing: a streamlined, ultra-simple and fast library preparation protocol for strand specific mrna library construction. Front. plant science 6 (2015).

34. AM Bolger, M Lohse, B Usadel, Trimmomatic: a flexible trimmer for illumina sequence data. Bioinformatics 30, 2114–2120 (2014).

35. NL Bray, H Pimentel, P Melsted, L Pachter, Near-optimal probabilistic rna-seq quantification. Nat. biotechnology 34, 525 (2016).

36. MD Robinson, DJ McCarthy, GK Smyth, edger: a bioconductor package for differential expression analysis of digital gene expression data. Bioinformatics 26, 139–140 (2010).

37. JA Romero Navarro, et al., A study of allelic diversity underlying flowering-time adaptation in maize landraces. Nat. genetics 49, 476 (2017).

38. DE Runcie, L Crawford, Fast and flexible linear mixed models for genome-wide genetics. PLOS Genet. 15, 1–24 (2019).

39. SC Potter, et al., HMMER web server: 2018 update (2018).

40. S El-Gebali, et al., The pfam protein families database in 2019. Nucleic acids research 47, D427–D432 (2019).

41. E Cimen, SE Jensen, ES Buckler, Building a tRNA thermometer to estimate microbial adaptation to temperature. Nucleic Acids Res. (2020).

42. SR Eddy, Accelerated profile HMM searches. PLoS computational biology 7, e1002195 (2011).

43. Z Li, J Tang, F Guo, Identification of 14-3-3 proteins Phosphopeptide-Binding specificity using an Affinity-Based computational approach. PloS one 11, e0147467 (2016).

44. N Patterson, et al., Ancient admixture in human history. Genetics 192, 1065–1093 (2012).

45. SH Martin, JW Davey, CD Jiggins, Evaluating the use of ABBA-BABA statistics to locate introgressed loci. Mol. biology evolution 32, 244–257 (2015).

46. E Gonzalez-Segovia, et al., Characterization of introgression from the teosinte zea mays ssp. mexicana to mexican highland maize. PeerJ 7, e6815 (2019).

47. R Bukowski, et al., Construction of the third generation zea mays haplotype map. GigaScience (2017).

48. E Paradis, pegas: an R package for population genetics with an integrated modular approach. Bioinformatics 26, 419–420 (2010).

49. SA Flint-Garcia, et al., Maize association population: a high-resolution platform for quantitative trait locus dissection: High-resolution maize association population. The Plant journal: for cell molecular biology 44, 1054–1064 (2005).

50. KAG Kremling, et al., Dysregulation of expression correlates with rare-allele burden and fitness loss in maize. Nature 555, 520–523 (2018).

51. HY Hung, et al., ZmCCT and the genetic basis of day-length adaptation underlying the postdomestication spread of maize (2012).

52. SC Stelpflug, et al., An Expanded Maize Gene Expression Atlas based on RNA Sequencing and its Use to Explore Root Development. The plant genome 9, 0 (2016).

53. AJ Waters, et al., Natural variation for gene expression responses to abiotic stress in maize. The Plant J. 89, 706–717 (2017).

54. WA Ricci, et al., Widespread long-range cis-regulatory elements in the maize genome. Nat. plants 5, 1237–1249 (2019).

55. O Trott, AJ Olson, AutoDock vina: improving the speed and accuracy of docking with a new scoring function, efficient optimization, and multithreading. J. computational chemistry 31, 455–461 (2010).

56. M Baek, et al., Accurate prediction of protein structures and interactions using a three-track neural network. Science 373, 871–876 (2021).

57. Y Nakamura, et al., High-Resolution crystal structure of arabidopsis FLOWERING LOCUS T illuminates its Phospholipid-Binding site in flowering. iScience 21, 577–586 (2019).

58. MW Maciejewski, et al., Nmrbox: A resource for biomolecular nmr computation. Biophys. J. 112, 1529–1534 (2017).

