## Supplementary File 8 for "An adaptive teosinte *mexicana* introgression modulates phosphatidylcholine levels and is associated with maize flowering time"

**Fragments of GRMZM2G353444 (B73) gene**

**Fragment 1**

**AACTTGTGTGATGAAGGTTCCACCG**TTCCAGTGTAACCACAGCATGCTCAACTCTGCACTAATGGGTAGGGAACATATGCAGGATAAACGTGCGTCTAGGCTAGGAAGAGACTCTCACCTCGTCACCAGCGCAGTGAAAACGGCTATATATTTTACTTGACGGATGGTGTGGTGAGTGCGGGGTTACGTCCAAAAACCCATGCAGAGCTATGATTTTATTTATTTTTTCGTCACAGGGTTCAAATGCCACATCAATATAATTAGCTCTAGCTAGCTCGGCTCGTCACCGTCACCGTCACAGTGCACATCTTGGACTAGCTCTTTGGTTGTGGTTGTGGCGATGGAGAATTGGAGACACCGGGCCGGCCCCGCTTCGATGCGTGGATCACGTGTGTGTCGATCGGTACCACAAAACACATGTCCTTGGTTTCTTGCGTTGGGCGGGGCCCGCACAACGCGTTACGCATGCATGCATGCATGTAGCGTTTGCTGTATTGCTGTGCTGTGTTACCATTGGCCGCGCGACATTGCTCCACGACGACTAGGGACCACCCAGACCCACCAAGG**CCGCAAGGTAGGATATTTCATGGATTATTATAAGAATTTAGCGCATCTTGGTACGCTTCCCGTTCAAGTGTTCTTTTGAAAAGAAAGAAAAAAAAACTACGTACTCGAGGGGTCAGTTCTCATAGCTCTGGCTCGT**

**Fragment 2**

**CCGCAAGGTAGGATATTTCATGGATTATTATAAGAATTTAGCGCATCTTGGTACGCTTCCCGTTCAAGTGTTCTTTTGAAAAGAAAGAAAAAAAAACTACGTACTCGAGGGGTCAGTTCTCATAGCTCTGGCTCGT**AGCCTGACCTGAAAGCAACTCCGGCTAGCTCCTCTATCAATCACTATAAAAAACGATTCCTCCTCCTAAAATATCTGATATACGGCTCTCTCGTAGAAGATGTTGCTAGACGCTAGAGTCATAGCCAGAACAGTTTTGAGCAGGACCTTAGAAAGCTAACGGTCGAGCTTAGCTTAGCTTAGCTCAGTTGGTTGGGTGAAACCTGACTAGTCGAGCACTTTCCAGTGCTCGAGTGTACGTACGTACGTACGTGTGCTCATATTATTCATATATTTGTTCTAGAATTTAACAACGTTATTGTTTTTTGTGATAGGTGGCGTATTTATCCGTTGAGTGTACGTAAGTCTCAATTTATAATTTGTTTGCCTTTTTTTTACTAAATTGGATCCACTCATCTTATTAATTTTTTTTGCGAAAAATGAAAAAATCAAAGTCATACTTAAAGTATTTTATATGCTAAATGACATCTCAATAAAAACTAATAATTATTTTTTAATAA**TACGAGCTGGTTAAACTTGGGGATAAAAATTCAAATGGATTATAAATTGAAACGAAGGGATCTCATTTAATATGTGGTATCTTAAGAGGTCTTTAGGGCCTTAATTGAAAAAAATGCAAGAGACAGACTCTTCTACACGA**

**Fragment 3**

**TACGAGCTGGTTAAACTTGGGGATAAAAATTCAAATGGATTATAAATTGAAACGAAGGGATCTCATTTAATATGTGGTATCTTAAGAGGTCTTTAGGGCCTTAATTGAAAAAAATGCAAGAGACAGACTCTTCTACACGA**CTCTCCATACATATTATCTCTTAGCTACATTTATGATCTGTAAGCTCAGGGGTGTATCATGCAGTCTCTTTGTTGTCTCCTATACTTAGAAAACCGATCCGAAGTCTTAGAGTTGTACATGCCCCTTAGAAGGCTGTTACATTATTTTGAGTTAGTGTGGATAATATATGATATAGAAATGACACAGTGTTTGCGTGTTTTTATAGAGGTGATATTTAGAGTGCCGACTTTGTTAGTGTTTTGAGTTAGCGAGGATAATGTGATATATAGAATATTTCAACTGAGATATTTGTATGGGCCGGTTCCGCACGCGGTGCACGCGCCCAAGTCCCCAACCATTGGTACCGATAGGAGTATAGCTAGGACTGGACGGTGGTCCCATCCACACATGGGCAGTACAGAAGCCATGTTGCACCATCAAGGGACGAGCTGCATTTTCACGTGGAGTCGTGGACGCATTATCATGTGGCTAGCTGTGCCCTGTGGGCCGCGGCGCCAAGACATGCCAAAGG**CCCGTCGTACTCGTGCTACTGCATGTTGAGCCACTTGTCAGTTCTCATCCCATGCACGCATCGCCATATATCAATGGCGAGCTCCTCCTCCTCCGAGCACCGAGCTCAAACACACATCAAGAAAGCA**

**Fragment 4**

**CCCGTCGTACTCGTGCTACTGCATGTTGAGCCACTTGTCAGTTCTCATCCCATGCACGCATCGCCATATATCAATGGCGAGCTCCTCCTCCTCCGAGCACCGAGCTCAAACACACATCAAGAAAGCA**AGAAAGATAAGACTGTAGCTAGCCACCGCGCGCACGAGTAGTCTAGCTAGACTACACTCACATTCAAAACAAACACACAGCGGCGTGCTGCGTGATCTCTGACCGCGTCGACAACTACGGTGACCGCAGCACCAAGCAGCCGATGGCGGCCACCGTATCTTCCTGCCTGAGCCTCGTCCCCACCGTGCACCACCGGGCCGGCGGGTTCCCCGCTGTGGCACTGACGAACACGCTTCAGCGCCGTCGTCGTTCTCGCCGCTATCTCGTCGTCGTCGCCTCGGCGACGAGAGCCGAACAGGAGGCCGCCCCTGTGACGATCGAGGACGTGGACCTCAGCTCTCATCAGGCGGCGCCCGACGATGGGGAGCTCGCCGCGCGGTGGCCGGAGATCCACGGCAGCAACAACTGGGAGGGCCTGCTGGACCCCATCGACGGCGTGCTCCTCCAGGAGCTCATCCGCTACGGCGAGTTCGCCCAGGCCACCTACGACAGCTTCGACTACGACCGCTTCTCCCCCTACTGCGGCA**GCTGCAAGTACCCGGCGAAGACCTTCTTCCACGACGTCGGCCTCGGCGGCATCGGCTACGAGGTCACCCGCTACCTCTACGCCACCTGCAACGACCTCAAGTTCCCCAACT**

**Fragment 5**

**GCTGCAAGTACCCGGCGAAGACCTTCTTCCACGACGTCGGCCTCGGCGGCATCGGCTACGAGGTCACCCGCTACCTCTACGCCACCTGCAACGACCTCAAGTTCCCCAACT**TCGGCATCAAGACGGCCGCCAACGCCAAGATGTGGAGCGAGTCCGGCACGTTCATCGGGTACGTGGCCGTGTCCACCGACGAGGAGACGGCGCGGCTCGGCCGCCGGGACATCGCCGTGGCGTGGCGCGGCACCATCACCCGCCTCGAGTGGGTGGCCGATCTGACGGCCAACCAGATCCCCCTGCGCGAGACGGGCGTCCCCTGCCCGGACCCCGACGTGAAGGTGGAGCGAGGGTTCGTGGCGCTGTACACGGACAAGGGCACCGGCTGCCGCTTCTGCCGGTACTCGGCGCGGGAGCAGGTGCTGGCCGAGGTGCGCAAGCTGGTGGATCTGTACCACGGGCGGGGCGAGCAGGTGAGCGTGACGGTCACGGGCCACAGCCTGGGCAGCGCGCTGGCGATGCTGTGCGCCTTCGACATCGCGGAGACGCGCGCGAACGTGTCGCCGGGCGACAGGGTGGCGCCGGTGTGCGTGTTCTCCTTCGCCGGCCCGCGCGTCGGGAACGTCGCCTTCCG**GAGGCGGTTCGAGCGGGAGCTGGGCGTGCGGGCGCTGCGCGTCGTGAACGTGCACGACAGCGTGCCCAAGGTGCCCGGCGTGTTCTTCAACGAGTCGGCGTTCCCG**

**Fragment 6**

**GAGGCGGTTCGAGCGGGAGCTGGGCGTGCGGGCGCTGCGCGTCGTGAACGTGCACGACAGCGTG
CCCAAGGTGCCCGGCGTGTTCTTCAACGAGTCGGCGTTCCCG**GAGCTCGTCCTGCGGGCGGCGGACAGGCTGGGCCTCGGCGGCGTGTACACGCACCTGGGCGTGCTGCTGCAGCTGGACCACAAGGTGTCGCCGTTCCTCAAGGAGACCCTGGACCTCTCGTGCTACCACAACCTGGAGGCGCACCTGCACCTGCTGGACGGCTTCCGCGGCTCCGGCGCGGGGTTCGAGCCCCGCGGGAGGGACCCCGCGCTGGTGAACAAGTCCACCGACTTCCTCCGGGAGGACCACATGGTGCCGCCCGTGTGGTACCAGGCGGAGAACAAGGGCATGGTGAGGACGGAGGACGGCCGGTGGGTGCTGCCGCCGCGGCAGAGGGTGCTCGACGACCACCCGGAGGACACTGACCATCACCTCCAGCGGCTCGGTCTCACCGCGTGAAGCTCAGCAATACTCTGATAGTAGCGTTAATTAGTGCGTGTGGCGTTCGCGTACGTGCGCGGGGCAGTCGGTTACTCTGTTGCCGCGCGCGGCAGCTCAGTGGCGCGTTCATGCCGTGCATGATGCGGGTGACGTGCGGGATGGGCACGTCGCTGCCTGAGAAGCTCGCACCGTTGGCGTTCGTGTGGCTGCAGAAAGGTCGTCTTGGATTGTAATTCTCTCTACTTATTTTTGTAAATTGAC**CCAGAAAAGCTGCTGCTTGCAGCGCC**

**AACTTGTGTGATGAAGGTTCCACCG**TTCCAGTGTAACCACAGCATGCTCAACTCTGCACTAATGGGTAGGGAACATATGCAGGATAAACGTGCGTCTAGGCTAGGAAGAGACTCTCACCTCGTCACCAGCGCAGTGAAAACGGCTATATATTTTACTTGACGGATGGTGTGGTGAGTGCGGGGTTACGTCCAAAAACCCATGCAGAGCTATGATTTTATTTATTTTTTCGTCACAGGGTTCAAATGCCACATCAATATAATTAGCTCTAGCTAGCTCGGCTCGTCACCGTCACCGTCACAGTGCACATCTTGGACTAGCTCTTTGGTTGTGGTTGTGGCGATGGAGAATTGGAGACACCGGGCCGGCCCCGCTTCGATGCGTGGATCACGTGTGTGTCGATCGGTACCACAAAACACATGTCCTTGGTTTCTTGCGTTGGGCGGGGCCCGCACAACGCGTTACGCATGCATGCATGCATGTAGCGTTTGCTGTATTGCTGTGCTGTGTTACCATTGGCCGCGCGACATTGCTCCACGACGACTAGGGACCACCCAGACCCACCAAGG**CCGCAAGGTAGGATATTTCATGG**ATTATTATAAGAATTTAGCGCATCTTGGTACGCTTCCCGTTCAAGTGTTCTTTTGAAAAGAAAGAAAAAAAAACTACGTACTCGAGG**GGTCAGTTCTCATAGCTCTGGCTCGT**AGCCTGACCTGAAAGCAACTCCGGCTAGCTCCTCTATCAATCACTATAAAAAACGATTCCTCCTCCTAAAATATCTGATATACGGCTCTCTCGTAGAAGATGTTGCTAGACGCTAGAGTCATAGCCAGAACAGTTTTGAGCAGGACCTTAGAAAGCTAACGGTCGAGCTTAGCTTAGCTTAGCTCAGTTGGTTGGGTGAAACCTGACTAGTCGAGCACTTTCCAGTGCTCGAGTGTACGTACGTACGTACGTGTGCTCATATTATTCATATATTTGTTCTAGAATTTAACAACGTTATTGTTTTTTGTGATAGGTGGCGTATTTATCCGTTGAGTGTACGTAAGTCTCAATTTATAATTTGTTTGCCTTTTTTTTACTAAATTGGATCCACTCATCTTATTAATTTTTTTTGCGAAAAATGAAAAAATCAAAGTCATACTTAAAGTATTTTATATGCTAAATGACATCTCAATAAAAACTAATAATTATTTTTTAATAA**TACGAGCTGGTTAAACTTGGGGATA**AAAATTCAAATGGATTATAAATTGAAACGAAGGGATCTCATTTAATATGTGGTATCTTAAGAGGTCTTTAGGGCCTTAATTGAAAAAAA**TGCAAGAGACAGACTCTTCTACACGA**CTCTCCATACATATTATCTCTTAGCTACATTTATGATCTGTAAGCTCAGGGGTGTATCATGCAGTCTCTTTGTTGTCTCCTATACTTAGAAAACCGATCCGAAGTCTTAGAGTTGTACATGCCCCTTAGAAGGCTGTTACATTATTTTGAGTTAGTGTGGATAATATATGATATAGAAATGACACAGTGTTTGCGTGTTTTTATAGAGGTGATATTTAGAGTGCCGACTTTGTTAGTGTTTTGAGTTAGCGAGGATAA
TGTGATATATAGAATATTTCAACTGAGATATTTGTATGGGCCGGTTCCGCACGCGGTGCACGCGCCCAAGTCCCCAACCATTGGTACCGATAGGAGTATAGCTAGGACTGGACGGTGGTCCCATCCACACATGGGCAGTACAGAAGCCATGTTGCACCATCAAGGGACGAGCTGCATTTTCACGTGGAGTCGTGGACGCATTATCATGTGGCTAGCTGTGCCCTGTGGGCCGCGGCGCCAAGACATGCCAAAGG**CCCGTCGTACTCGTGCTACTGCAT**GTTGAGCCACTTGTCAGTTCTCATCCCATGCACGCATCGCCATATATCAATGGCGAGCTCCTCCTCCTCCGAGCACCGA**GCTCAAACACACATCAAGAAAGCA**AGAAAGATAAGACTGTAGCTAGCCACCGCGCGCACGAGTAGTCTAGCTAGACTACACTCACATTCAAAACAAACACACAGCGGCGTGCTGCGTGATCTCTGACCGCGTCGACAACTACGGTGACCGCAGCACCAAGCAGCCGATGGCGGCCACCGTATCTTCCTGCCTGAGCCTCGTCCCCACCGTGCACCACCGGGCCGGCGGGTTCCCCGCTGTGGCACTGACGAACACGCTTCAGCGCCGTCGTCGTTCTCGCCGCTATCTCGTCGTCGTCGCCTCGGCGACGAGAGCCGAACAGGAGGCCGCCCCTGTGACGATCGAGGACGTGGACCTCAGCTCTCATCAGGCGGCGCCCGACGATGGGGAGCTCGCCGCGCGGTGGCCGGAGATCCACGGCAGCAACAACTGGGAGGGCCTGCTGGACCCCATCGACGGCGTGCTCCTCCAGGAGCTCATCCGCTACGGCGAGTTCGCCCAGGCCACCTACGACAGCTTCGACTACGACCGCTTCTCCCCCTACTGCGGCA**GCTGCAAGTACCCGGCGAAGACCTTC**TTCCACGACGTCGGCCTCGGCGGCATCGGCTACGAGGTCACCCGCTACCTCTACGCCACCT**GCAACGACCTCAAGTTCCCCAACT**TCGGCATCAAGACGGCCGCCAACGCCAAGATGTGGAGCGAGTCCGGCACGTTCATCGGGTACGTGGCCGTGTCCACCGACGAGGAGACGGCGCGGCTCGGCCGCCGGGACATCGCCGTGGCGTGGCGCGGCACCATCACCCGCCTCGAGTGGGTGGCCGATCTGACGGCCAACCAGATCCCCCTGCGCGAGACGGGCGTCCCCTGCCCGGACCCCGACGTGAAGGTGGAGCGAGGGTTCGTGGCGCTGTACACGGACAAGGGCACCGGCTGCCGCTTCTGCCGGTACTCGGCGCGGGAGCAGGTGCTGGCCGAGGTGCGCAAGCTGGTGGATCTGTACCACGGGCGGGGCGAGCAGGTGAGCGTGACGGTCACGGGCCACAGCCTGGGCAGCGCGCTGGCGATGCTGTGCGCCTTCGACATCGCGGAGACGCGCGCGAACGTGTCGCCGGGCGACAGGGTGGCGCCGGTGTGCGTGTTCTCCTTCGCCGGCCCGCGCGTCGGGAACGTCGCCTTCCG**GAGGCGGTTCGAGCGGGAGCTGG**GCGTGCGGGCGCTGCGCGTCGTGAACGTGCACGACAGCGTG
CCCAAGGTGCCCGGCGTG**TTCTTCAACGAGTCGGCGTTCCCG**GAGCTCGTCCTGCGGGCGGCGGACAGGCTGGGCCTCGGCGGCGTGTACACGCACCTGGGCGTGCTGCTGCAGCTGGACCACAAGGTGTCGCCGTTCCTCAAGGAGACCCTGGACCTCTCGTGCTACCACAACCTGGAGGCGCACCTGCACCTGCTGGACGGCTTCCGCGGCTCCGGCGCGGGGTTCGAGCCCCGCGGGAGGGACCCCGCGCTGGTGAACAAGTCCACCGACTTCCTCCGGGAGGACCACATGGTGCCGCCCGTGTGGTACCAGGCGGAGAACAAGGGCATGGTGAGGACGGAGGACGGCCGGTGGGTGCTGCCGCCGCGGCAGAGGGTGCTCGACGACCACCCGGAGGACACTGACCATCACCTCCAGCGGCTCGGTCTCACCGCGTGAAGCTCAGCAATACTCTGATAGTAGCGTTAATTAGTGCGTGTGGCGTTCGCGTACGTGCGCGGGGCAGTCGGTTACTCTGTTGCCGCGCGCGGCAGCTCAGTGGCGCGTTCATGCCGTGCATGATGCGGGTGACGTGCGGGATGGGCACGTCGCTGCCTGAGAAGCTCGCACCGTTGGCGTTCGTGTGGCTGCAGAAAGGTCGTCTTGGATTGTAATTCTCTCTACTTATTTTTGTAAATTGAC**CCAGAAAAGCTGCTGCTTGCAGCGCC**AAAAAAAAAAACGATTTTCATCTTTCTGTTTGGTGTTTGCCATCGGACAACAGAGCGA
